# Laminin β4 is required for the development of human peripheral sensory neurons

**DOI:** 10.1101/2024.11.22.624899

**Authors:** Kenyi Saito-Diaz, Tripti Saini, Archie Jayesh Patel, Christina James, Kimata Safi Thomas, Trinity Nora Knight, Stephanie Beatrice Gogita, Nadja Zeltner

## Abstract

The extracellular matrix (ECM) is a mixture of glycoproteins and fibrous proteins that provide the biophysical properties necessary to maintain cellular homeostasis. ECM integrity is of particular importance during development, where it allows proper migration and cellular differentiation. Laminins are ECM heterotrimeric proteins consisting of α, β, and γ chains. There are five known α chains, four β chains, and three γ chains. Thus, there are 60 potential combinations for laminin trimers, however only 16 laminin trimers have been identified to date. Furthermore, none of them contain laminin β4 and its function is unknown. Here, we sought to characterize the role of *LAMB4* (the gene encoding laminin β4) during human embryonic development of the peripheral sensory nervous system. Using human pluripotent stem cells (hPSCs), we found that *LAMB4* is expressed in the ectoderm in the early stages of sensory neuron (SN) specification. SNs, part of the peripheral nervous system, are specialized neurons that detect pain, temperature, and touch. Surprisingly, more than 20 million people in the US have some form of peripheral nerve damage (including SNs), however there are very few treatment options available. Learning about the biology of peripheral neurons will uncover potential new therapeutic targets, thus we focused on understanding the effects of *LAMB4* in SNs. First, we knocked out *LAMB4* in hPSCs, using CRISPR/Cas9, and found that loss of *LAMB4* impairs the migration of the SN progenitors neural crest cells (NCCs) and harms SN development and survival. To assess if *LAMB4* has clinical relevance, we studied the genetic disorder Familial Dysautonomia (FD), which specifically affects the peripheral nervous system. FD is caused by a mutation in *ELP1* (a component of the Elongator complex) leading to developmental and degenerative defects in SNs. A previous report showed that patients with severe FD harbor additional single nucleotide variants in *LAMB4*. We found that these variants sharply downregulate the expression of *LAMB4* and laminin β4 levels in SNs differentiated from induced pluripotent stem cells (iPSCs) reprogramed from patients with severe FD. Moreover, a healthy ECM is sufficient to rescue the developmental phenotypes of FD, further confirming that ECM defects contribute significantly to the etiology of FD. Finally, we found that *LAMB4*/laminin β4 is necessary for actin filament accumulation and it interacts with laminin α4 and laminin γ3, forming the laminin-443, a previously unreported laminin trimer. Together, these results show that *LAMB4* is a critical, but largely unknown gene required for SN development and survival.

## Introduction

The extracellular matrix (ECM) is a dynamic network of proteins, glycoproteins, and proteoglycans that act as a cellular scaffold, providing the ideal environment to promote and maintain cellular homeostasis^1,2^. The ECM is a very dynamic structure, and it is constantly remodeled to support cell homeostasis^3^ and provides biophysical and biochemical cues required for cellular migration, differentiation, and survival during development^4^. For instance, neural crest cells (NCCs) arise from the border of the neural plate and the non-neural ectoderm of the developing embryo^5,6^. NCCs then migrate to different regions of the embryo in a process regulated by ECM remodeling and gradient of morphogens^7^, giving rise to sensory neurons (SNs) and autonomic neurons (part of the peripheral nervous system), glial cells, endocrine cells, craniofacial cartilage and bone, pigment cells, among others^8^. Changes in the biophysical properties of the ECM affect NCC differentiation^9^.

Laminins are major components of the ECM^10^. They are heterotrimeric proteins consisting of α, β, and γ chains that are expressed in different developmental stages for specific functions. For example, laminin-511 and laminin-111 are the most abundant laminins present during early development^11,12^, whereas laminin-523 is expressed only in the retinal outer membrane^13^. Laminins interact with the transmembrane proteins integrins^14^. They are heterodimer receptors that connect the ECM to intracellular components, resulting in activation of signaling pathways and reorganization of the cellular cytoskeleton^15^.

There are five α chains, four β chains, and three γ chains, which can assemble up to 60 different trimers, however, only 16 have been identified^15^. Of those, laminin β4 (expressed by the gene *LAMB4*) is understudied and no laminin trimer containing the laminin β4 chain has been identified. Laminin β4 downregulation has been linked to diverticulitis, a disease of the peripheral nervous system^16^. Additionally, single nucleotide variants of *LAMB4* have been identified in patients with severe symptoms of the genetic disease Familial dysautonomia (FD), which specifically affects the peripheral neurons. Thus, we hypothesized that *LAMB4*/laminin β4 expression is necessary for development and homeostasis of the peripheral nervous system and their progenitors, the NCCs.

To address this, we use human pluripotent stem cell (hPSC) technology, which allows us to study human development, including the study of cellular and molecular mechanisms in cells from all three germ layers endoderm, mesoderm, and ectoderm^17,18^. Additionally, cells obtained from patients can be reprogrammed into induced pluripotent stem cells (iPSCs), which contain the same genetic background as the originating patients and thus are invaluable for disease modeling^19^.

Here, we used hPSC technology and identified that *LAMB4* is expressed in the peripheral nervous system, and it is necessary for NCC migration, and development and survival of SNs. Patients with severe symptoms of the peripheral neuropathy FD harbor mutations in the *LAMB4* gene. We show that SNs differentiating from FD iPSCs express low levels of laminin β4. Additionally, we report that the ECM that is deposited by healthy cells is sufficient to rescue the developmental phenotypes observed in FD. Finally, we show that laminin β4 forms the laminin-443 and it is required for accumulation of actin filaments (F-actin) in SNs. Together, our results confirm that the ECM is critical for the development and function of SNs and that laminin β4 is required for SN development.

## Results

### Laminin β4 is expressed in early stages of sensory neuron differentiation

To understand the biological function of laminin β4 we sought to first assess the similarity between laminin β4 and the other laminin β chains. We started by asking whether metazoans express laminin β4. To do this, we compared the amino acid sequences of all the laminin β chains expressed in multiple species and generated a phylogenetic tree (**Fig. S1A**). We found that rodents (*M*. *musculus* and *R*. *norvegicus*) do not have a *LAMB4* ortholog, whereas other vertebrates do, such as frogs (*X*. *tropicalis*), zebrafish (*D*. r*erio*), dogs (*C*. *lupus*), and chickens (*G*. *gallus*, **Fig. S1A**). Our results also showed that laminin β4 is closely related to laminin β1 and β2 (**Fig. S1A**), and *LAMB4* is the ortholog expressed in the lowest number of the analyzed species: seven, compared to 11 for the other β chains. The lack of *LAMB4* in rodents, particularly in mice, could be one of the reasons why *LAMB4* has not been characterized yet and hints at its difficulty to study *in vivo*. We next asked whether laminin β4 shares similarities with other laminin β chains in humans. We didn’t find major differences between laminin β1, β2, and β4 chains, since they all shared the sequence of the N-terminal domain and 13 EGF-like domains (**Fig. 1A and S1B**). These domains are important for biological functions such as laminin network assembly, thus suggesting that they could bind similar proteins. In contrast, laminin β3 showed the shortest amino acid sequence and we only identified six EGF-like domains. On the C-terminal region, although all four laminin β chains showed similar length and number of domains, there were clear differences in the amino acid sequences (**Fig. 1A and S1C**). The C-terminal region bind to the α and γ chains, thus the differences between β chains might provide specificity during laminin assembly and be involved in the different affinities observed between laminin chains^20^. We next sought to understand which cell lineages express *LAMB4*. We first differentiated control hPSC-ctr-H9 cells into definitive endoderm (**Fig. S2A**). Although endoderm markers such as *SOX17*, *GATA4*, *GATA6*, and *FOXA2* were highly expressed, *LAMB4* was not (**Fig. S2B**). We also analyzed RNAseq of hindgut differentiated from hPSCs^21^, where we found that *LAMA1*, *LAMA5*, *LAMB1*, *LAMB2*, *LAMC1*, and *LAMC2* were highly expressed (green) but not *LAMB4* (red rectangle, **Fig. S2C**). Next, we assessed *LAMB4* throughout mesoderm and cardiomyocyte differentiation (**Fig. S2D**). During the differentiation we measured high expression of classic mesoderm (*TBXT*, *TBX6*, and *FOXF1*, **Fig. S2E**) and cardiomyocyte markers (*TNNT2* and *NKX2-5*, **Fig. S2F**), but not *LAMB4*. This was confirmed by analyzing published RNAseq data of human mesoderm and cardiomyocytes differentiated from hPSCs^22^. *LAMA1*, *LAMB1*, *LAMB2*, and *LAMC1* were highly expressed (green), but not *LAMB4* (red rectangle) during the assessed timepoints (**Fig. S2G**). Thus, we next investigated *LAMB4* expression in ectoderm, specifically in neural crest, by following a protocol to differentiate hPSCs into SNs using chemically defined conditions^23,24^ (**Fig. 1B**). In this protocol, SNs are differentiated by going through all the developmental stages observed *in vivo*^23^. We found that *LAMB4* mRNA is expressed in day 12 NCCs differentiated from hPSC-ctr-H9 cells, and it peaked in the early stages of SN specification, by around day 20 in our protocol (**Fig. 1C**). In contrast, laminin β4 isolated from cell lysates and the ECM increased over time and peaked at later stages of SN development (day 40-50, **Fig. 1D and E**). It is possible that this increase is caused by laminin β4 still being assembled and secreted to the ECM although not transcribed at high rates. To test this, we measured laminin β4 levels in the ECM alone. To do this, we lysed and removed the cells using ammonium hydroxide and resuspended the undisturbed ECM in Laemmli buffer followed by analysis by immunoblot^25^. Similar to our previous results, laminin β4 signal increased over time, which suggests that indeed laminin β4 was being continuously secreted (**Fig. 1F and G**). Finally, we confirmed our results by immunofluorescence, where we found that NCCs and SNs express laminin β4 (**Fig. 1H and I**). Together, our data shows that *LAMB4* is expressed in NCC lineages which include SNs.

**Figure 1.**
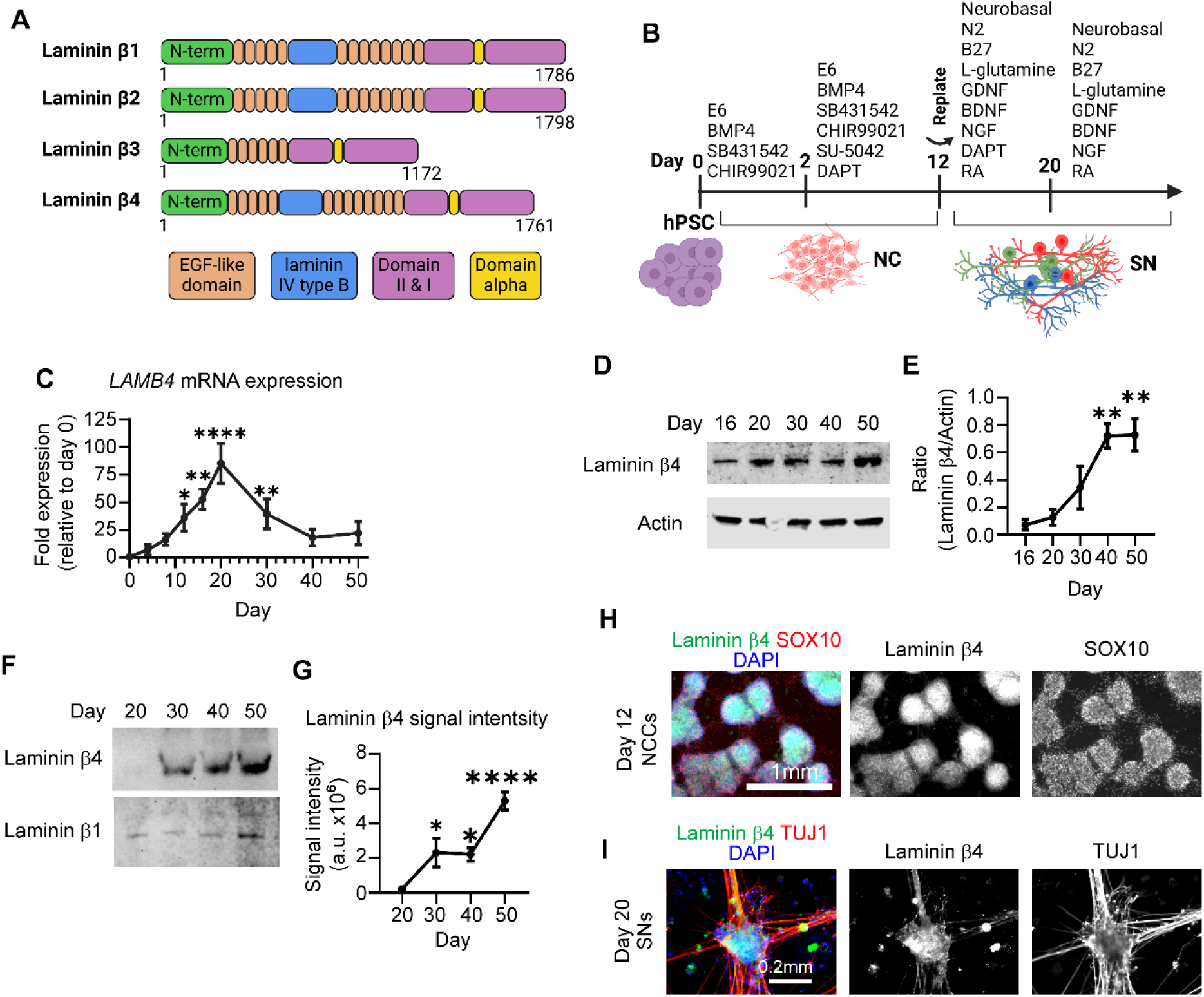
*LAMB4*/laminin β4 is expressed in neural crest cells (NCCs) and sensory neurons (SNs). **A)** Comparison of laminin β chains expressed in humans. **B)** Schematics of the NC and SN differentiation protocol. **C)** *LAMB4* expression during SN differentiation. hPSC-ctr-H9 SNs were harvested at the indicated times and mRNA expression of *LAMB4* was measured by RT-qPCR (n=3 biological replicates). **D)** Laminin β4 expression during SN development. Total protein was isolated from SNs differentiated from hPSC-ctr-H9 cells at the indicated times and immunoblotted for laminin β4 and actin. **E)** Signal intensity of immunoblots from **D)** was measured, quantified, and normalized to day 16 (n=3 biological replicates). **F)** Levels of laminin β4 in the ECM of SNs. hPSC-ctr-H9 SNs were harvested on the indicated days and the ECM was collected and immunoblotted for laminin β4. Plates were coated with laminin β1 and was used as a loading control. **G)** Signal intensity of immunoblots from **F)** was measured, quantified, and normalized to day 20 (n=4 biological replicates). **H)** Laminin β4 expression in NCCs. Day 12 NCCs differentiated from hPSC-ctr-H9 cells were fixed and stained for laminin β4, SOX10, and DAPI. **I)** Expression of laminin β4 in SNs. hPSC-ctr-H9 SNs were fixed on day 20 and stained for the laminin β4, TUJ1, and DAPI. For **C)**, **E),** and **G)**, one-way ANOVA followed by Dunnett’s multiple comparisons test. ns, non-significant, *p<0.05, **p<0.005, ****p<0.0001. Graphs show mean ± SEM.

### *LAMB4* is required for neural crest cell migration

Next, we asked what is the role *LAMB4* plays in SN development. To address this question, we knocked-out *LAMB4* in healthy hPSC-ctr-H9 cells using CRISPR/Cas9 (**Fig. S3A**). We identified a homozygous (*LAMB4^−/−^*) and a heterozygous (*LAMB4^+/−^*) clone by Sanger sequencing (**Fig. S3B**). Although there were no phenotypical differences in the hPSC colonies compared to the parental hPSC-ctr-H9 cell line (*LAMB4^+/+^*, **Fig. S3C**), laminin β4 levels at the mRNA and protein levels were reduced in *LAMB4^−/−^* and *LAMB4^+/−^* SNs (**Fig. S3D-F**). We then used these cell lines to ask whether *LAMB4* is required for the development of NCCs. We found that *LAMB4^−/−^* and *LAMB4^+/−^* can still differentiate into NCCs (**Fig. 2A**), however loss of *LAMB4* made the characteristic “ridges”, formed by accumulation of NCCs^22^, smaller by inspection by brightfield microscopy (**Fig. 2A, red arrows and B**). We first hypothesized that the reduced area was due to a decrease in the number of NCCs. We tested this by assessing the number SOX10^+^ NCCs by immunofluorescence. We indeed found a reduced number of large clusters of SOX10^+^ cells in the *LAMB4^−/−^* and *LAMB4^+/−^* cells, although there was a large number of single SOX10^+^ cells (**Fig. 2C**). To confirm our results, we quantified the number of cells expressing the migratory NCC marker CD49d (which correlates well with SOX10 at this developmental stage^23,26,27^) by flow cytometry analysis. There was no change in the number of CD49d^+^ cells in any of the lines (**Fig. 2D**), suggesting that the number of NCCs was not affected by *LAMB4*. During development, NCCs migrate and accumulate forming ganglia^28^. Because the number of NCCs did not change upon loss of *LAMB4*, but we saw a high number of individual SOX10^+^ cells by immunofluorescence (**Fig. 2C**), we hypothesized that loss of *LAMB4* impairs NCC migration. We first performed a scratch assay to test the migration of *LAMB4^+/+^*, *LAMB4^+/−^*, and *LAMB4^−/−^* NCCs. *LAMB4^+/−^* and *LAMB4^−/−^* NCCs failed to migrate after 48 hours compared to *LAMB4^+/+^* (**Fig. 2E and F**). To confirm this, we performed live-cell imaging to map the migration of NCCs. Agreeing with our previous results, *LAMB4^+/−^*, and *LAMB4^−/−^* cells migrated at a lower accumulated distance compared to the *LAMB4^+/+^* control (**Fig. 2G and H**). Finally, we characterized the expression of NCC genes in *LAMB4^+/−^* and *LAMB4^−/−^* NCCs. We found that *SOX10* expression was similar in all the cell lines (**Fig. 2I**), agreeing with our flow cytometry results (**Fig. 2D**). In contrast, expression of genes involved in SN specification such as *P75NTR*, *NGN1*, and *NGN2* was significantly downregulated in *LAMB4^+/−^* and *LAMB4^−/−^* NCCs (**Fig. 2I**). Together, our results show that *LAMB4* is necessary for NCC migration and for SN differentiation.

**Figure 2.**
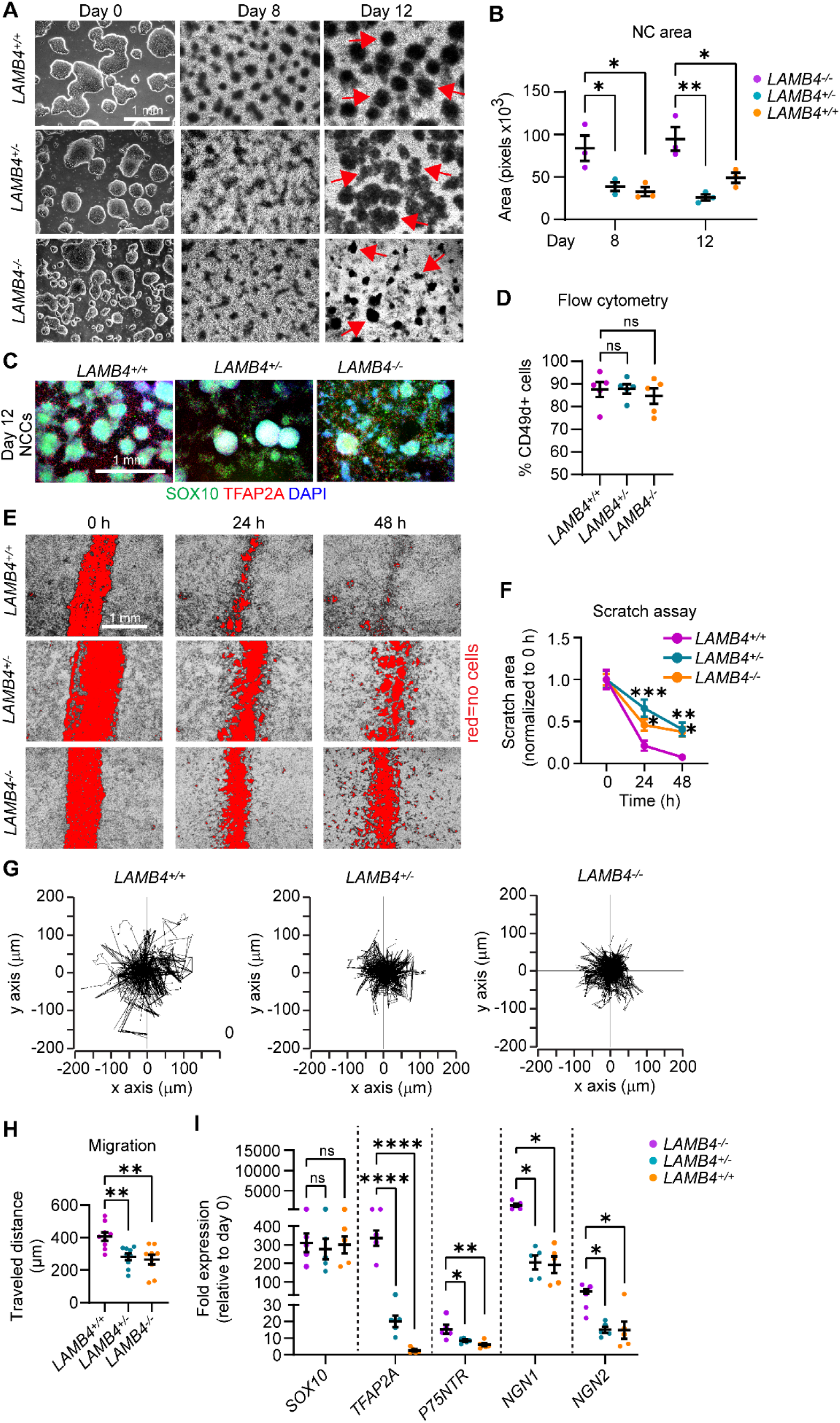
*LAMB4*/laminin β4 is required for NCC migration. **A)** Effects of *LAMB4* in NCCs. Brightfield images of colonies of *LAMB4^+/+^*, *LAMB4^+/−^*, and *LAMB4^−/−^* hPSCs and NCCs at different stages. Red arrows indicate “ridges” of NCCs. **B)** Quantification of the area of NCC-ridges in **A)**. Each dot indicates average of cell clusters that are part of the ridges (>70 clusters, n=3 biological replicates). **C)** Expression of NCC-markers upon loss of *LAMB4*. Representative immunofluorescence image of NCCs differentiated from *LAMB4^+/+^*, *LAMB4^+/−^*, and *LAMB4^−/−^* hPSCs were fixed and stained for SOX10 (green), TFAP2A (red), and DAPI (blue). **D)** Number of NCCs differentiated from *LAMB4* mutant hPSCs. NCCs differentiated from *LAMB4^+/+^*, *LAMB4^+/−^*, and *LAMB4^−/−^* cells were harvested, stained for the NCC surface marker CD49d, and analyzed using flow cytometry (n=5 biological replicates). **E)** Measurement of NCC migration by scratch assay. NCCs from *LAMB4* mutant cells were replated on day 8 and scratched when they reached confluency. Brightfield images were taken at 0, 24, and 48 hours after the scratch to follow NC migration into the scratched surface (shown in red). **F)** Scratched areas in **E)** (red) were measured and normalized to 0 hours. Average of 5-10 wells per condition are plotted (n=3 biological replicates) **G)** Migration of NCCs by live-cell imaging. Day 8 NCCs from *LAMB4^+/+^*, *LAMB4^+/−^*, and *LAMB4^−/−^* cells were replated and imaged every 10 minutes for 18 hours. Individual cells were tracked, and their traveled distance and direction were measured and plotted. **H)** Accumulated distance traveled by NCCs measured in **G)**. The average of 9 wells per condition are plotted (n=3 biological replicates). **I)** Expression of NCC-related genes upon loss of *LAMB4*. RNA from *LAMB4^+/+^*, *LAMB4^+/−^*, and *LAMB4^−/−^* NCCs (day 12) was isolated, and mRNA levels were measured using RT-qPCR (n=5 biological replicates). For **B)**, **D)**, **H),** one-way ANOVA followed by Tukey’s multiple comparisons test. For **I)**, one-way ANOVA followed by Dunnett’s multiple comparisons test. For **F)**, two-way ANOVA followed by Tukey’s multiple comparisons test. ns, non-significant, *p<0.05, **p<0.005, ****p<0.0001. Graphs show mean ± SEM.

### *LAMB4* is required for the development of sensory neurons

Our results suggest that *LAMB4* plays an important role in directing NCCs into a SN fate. Thus, we asked whether loss of *LAMB4* negatively affects the development of SNs in our human *in vitro* system. We found that on day 20 of our differentiation protocol, the number of neurons differentiated from *LAMB4^+/−^* and *LAMB4^−/−^* hPSCs was decreased (**Fig. 3A**). Moreover, by day 50, the size of the SN clusters, reminiscent to the ganglia observed *in vivo*^29^, were reduced in *LAMB4^+/−^* SNs and virtually inexistent in *LAMB4^−/−^* SNs (**Fig. 3A**). We confirmed these results by immunofluorescence, where the clusters of SNs, stained for the SN marker BRN3A and the pan-neuronal marker TUJ1, were reduced in *LAMB4^+/−^* and *LAMB4^−/−^* lines compared to the parental control (**Fig. 3B and C**). Additionally, the size of clusters and the number of SNs stained for ISL1^+^ (SN marker) and PRPH^+^ (peripheral neuron marker) were also reduced (**Fig. 3B and D**). When we quantified the number of SNs by flow cytometry we found that, when compared to wild type, both *LAMB4^+/−^* and *LAMB4^−/−^* were reduced, *LAMB4* heterozygous hPSCs differentiate more efficiently than homozygous *LAMB4* knockout cells (*LAMB4^−/−^*), suggesting that the *LAMB4* expression level is important for SN development (**Fig. 3E**). These results showed that loss of *LAMB4* impaired the development of SNs *in vitro* without changing the number of NCCs. We also found an increase in the number of non-neuronal cells expressing alpha-smooth muscle actin (αSMA) differentiated from *LAMB4^+/−^* and *LAMB4^−/−^* hPSCs compared to the parental control (**Fig. S4A and B**). Furthermore, *ACTA2*, the gene expressing αSMA, was upregulated in *LAMB4^+/−^* and *LAMB4^−/−^* cells (**Fig. S4C**). On the other hand, we didn’t see upregulation of genes expressed by sympathetic neurons (*ASCL1*), motor neurons (*MNX1*), enteric neurons (*EDRNB*), and other CNS cells (*OLIG2*, **Fig. S4C**). Together our results show that in the absence of *LAMB4*, NCCs do not differentiate into SNs efficiently and the number non-neuronal αSMA^+^ cells increases. These results strengthen our hypothesis that *LAMB4* is necessary for NCC and SN-specification, and in its absence, NCCs take a non-neuronal cell fate.

**Figure 3.**
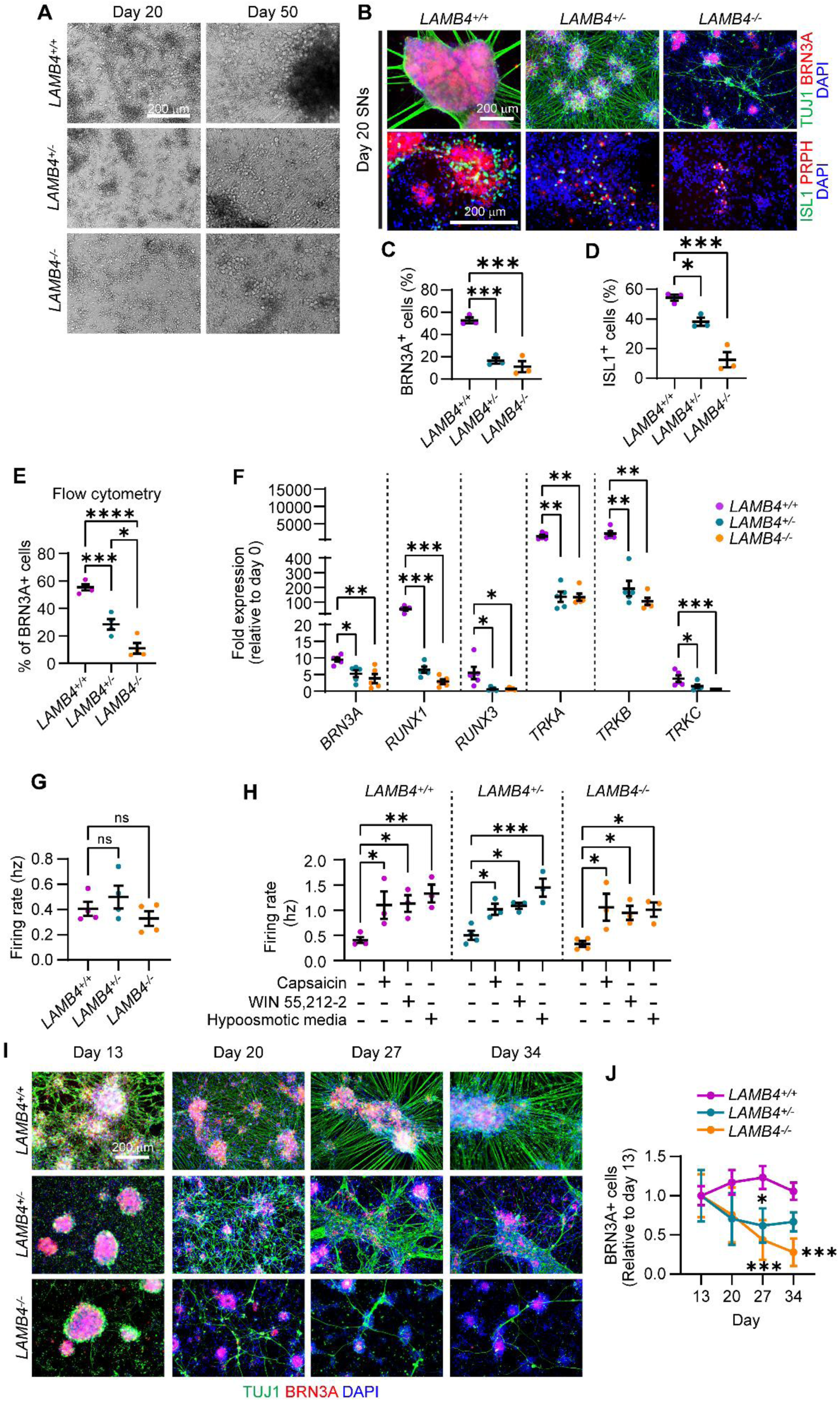
*LAMB4* is necessary for SN development and survival. **A)** Effects of LAMB4 downregulation in SNs. Brightfield images of SNs differentiated from *LAMB4^+/+^*, *LAMB4^+/−^*, and *LAMB4^−/−^* hPSCs at indicated days. **B)** Expression of SN markers upon loss of *LAMB4*. *LAMB4^+/+^*, *LAMB4^+/−^*, and *LAMB4^−/−^* SNs were fixed on day 20 and stained for peripheral neuron markers (TUJ1 and PRPH) and SN markers (BRN3A, ISL1). Nuclei were stained with DAPI. **C)** Percentage of BRN3A^+^ cells in **B)**. Normalized to DAPI. **D)** Percentage of ISL1^+^ cells in **B)**. Normalized to DAPI. **E)** Quantification of the number of SNs differentiated from *LAMB4* mutant hPSCs. SNs from *LAMB4^+/+^*, *LAMB4^+/−^*, and *LAMB4^−/−^* cells were harvested on day 20. SNs were then fixed, stained for BRN3A, and analyzed by flow cytometry (n=4 biological replicates). **F)** Expression of SN markers in *LAMB4* mutant SNs. RNA from *LAMB4^+/+^*, *LAMB4^+/−^*, and *LAMB4^−/−^* SNs was isolated on day 20 and gene expression was measured by RT-qPCR (n=5 biological replicates). **G)** Electrical activity of SNs upon loss of *LAMB4*. The firing rate of *LAMB4^+/+^*, *LAMB4^+/−^*, and *LAMB4^−/−^* SNs was measured using multi-electrode array (MEA). Each dot represents the mean firing rate of 6 wells measured over 40 days (n=4 biological replicates). **H)** Electrophysiological changes of SNs to pharmacological nociceptor or mechanoreceptor activators. *LAMB4*-mutant SNs were incubated with nociceptor agonists (0.25 µM capsaicin and 1 µM WIN55,212-2) and a mechanoreceptor activator (hypoosmotic medium) and the electrical activity was measured using MEA. Each dot represents the mean firing rate of 6 wells measured over 40 days (n=4 biological replicates). **I)** Degeneration of *LAMB4* mutant SNs. *LAMB4^+/+^*, *LAMB4^+/−^*, and *LAMB4^−/−^* SNs were cultured in plates coated with fibronectin and poly-L-ornithine and reduced NGF concentration (1 ng/mL) and fixed on the indicated days. Cells were then stained for the neuronal marker TUJ1, the SN marker BRN3A, and DAPI. **J)** Quantification of BRN3A^+^ SNs from **I)** (n=3-4 biological replicates). For **C)**, **D)**, **E)**, and **G)** one-way ANOVA followed by Tukey’s multiple comparisons test. For **F)** and **H)**, one-way ANOVA followed by Dunnett’s multiple comparisons test. For **J)**, two-way ANOVA followed by Šídák’s multiple comparisons test. ns, non-significant, *p<0.05, **p<0.005, ***p<0.001, ****p<0.0001. Graphs show mean ± SEM.

The three main subtypes of SNs found in the human dorsal root ganglia are nociceptors, mechanoreceptors, and proprioceptors, which detect pain, touch, and body position relative to space, respectively^28^. Since *LAMB4* is necessary for SN development, we next asked whether its loss impacts a particular subtype or all of these SN types. During development, different genes are expressed to promote specification of SNs to a unique subtype. All SNs express *BRN3A*, whereas *RUNX1* and *TRKA* are expressed by nociceptors during development. In contrast, mechanoreceptors express *TRKB*, and proprioceptors express *TRKC*. Additionally, progenitors of both mechanoreceptors and proprioceptors express *RUNX3*^28^. We first analyzed the expression of these genes by RT-qPCR. *BRN3A* was downregulated in *LAMB4^+/−^* and *LAMB4^−/−^* SNs compared to *LAMB4^+/+^* cells (**Fig. 3F**). Moreover, the nociceptor-related genes *RUNX1* and *TRKA* were also downregulated, as well as *RUNX3*, *TRKB*, and *TRKC*, which are expressed by mechanoreceptors and proprioceptors (**Fig. 3F**). Additionally, the number of cells expressing TRKA, TRKB, and TRKC was reduced in *LAMB4* mutant SNs measured by flow cytometry (**Fig. S4D**). Thus, *LAMB4* is required for the development of all the three main SN subtypes. We next tested whether the neurons that developed were electrically active. We didn’t find any difference in the firing rate between *LAMB4^+/+^, LAMB4^+/−^*, and *LAMB4^−/−^* SNs (**Fig. 3G**). Agreeing with this, the number, duration, frequency, and intervals of bursts were the same among the three lines (**Fig. S4E-H**). Furthermore, when we activated nociceptors with the agonists capsaicin and WIN55,212-2^23^ the firing rate of *LAMB4^+/+^, LAMB4^+/−^*, and *LAMB4^−/−^* SNs was similarly increased (**Fig. 3F**). This was also observed when *LAMB4^+/+^, LAMB4^+/−^*, and *LAMB4^−/−^* mechanoreceptors were activated using hypoosmotic medium^23^ (**Fig. 3H**), suggesting that *LAMB4* does not play a role in SN function.

The ECM plays important roles in the homeostasis of different cell types, including neurons^30^. Thus, we asked whether *LAMB4* is necessary for the survival of SNs. Our differentiation protocol had been optimized to assure the development and survival of wild type SNs^31^. To assess degeneration in non-wild type SNs, we first developed a modified protocol that accelerates degeneration in more vulnerable (for example diseased) lines, while still remaining robust cell survival in healthy SNs. This protocol consists of: (1) reducing the concentration of nerve growth factor (NGF) in the differentiation medium and (2) lack of coated laminin in the plates during the differentiation^31^. With this approach, we found that *LAMB4^+/−^* and *LAMB4^−/−^* SNs degenerate faster compared to *LAMB4^+/+^* (**Fig. 3I and J**). However, *LAMB4^−/−^* SNs die at a faster rate compared to *LAMB4^+/−^*. Together, our results show that *LAMB4* is required for both development and survival, but not function of SNs. Next, we sought to test whether altered *LAMB4* expression has clinical implications.

### A healthy extracellular matrix rescues the developmental phenotypes of the peripheral neuropathy Familial Dysautonomia

The peripheral neuropathy Familial Dysautonomia (FD) is a devastating genetic disease that specifically targets peripheral neurons^32^. It is caused by a mutation in the elongator complex scaffold protein *ELP1*^33,34^. FD was one of the first diseases to be modeled using the iPSC technology^35^. While 99.5% of all FD patients harbor the precise *ELP1* mutation, clinical symptoms vary widely among patients. The reasons for this discrepancy were unclear. To address this question, we reported that the severity of FD symptoms can be recapitulated in our *in vitro* system^36^. SNs differentiated from iPSCs of patients with mild symptoms showed only degenerative, but not developmental phenotypes. In contrast, iPSCs reprogrammed from patients with severe symptoms showed significant neurodevelopmental impairment, as well as neurodegeneration^36^ (**Fig. 4A**). We found that iPSCs from severe FD, but not mild FD patients harbored variants in *LAMB4,* which could account for the phenotypical differences^36^. Thus, we focused on addressing whether these variants in the FD patient iPSC lines affect *LAMB4* expression. The ECM is important to maintain the availability of growth factors required for neuron development^37^. Therefore, loss of *LAMB4* could affect the biophysical properties of the ECM and change the diffusion and availability of growth factors and signaling molecules. To confirm whether changes in the ECM affect the development of SNs in iPSCs derived from FD patients, we isolated the ECM of healthy cells as previously reported^25^. First, we differentiated healthy hPSC-ctr-H9 cells into NCCs and SNs, followed by lysis of the cells to maintain the healthy ECM. Severe FD iPSCs (iPSC-FD-S3) were then differentiated on top of the isolated healthy ECM (**Fig. 4B**). ECM from hPSC-ctr-H9 NCCs was sufficient to increase the area of the SOX10^+^ ridges and the number of NCCs (**Fig. 4C and D**). We observed the same results when iPSC-FD-S3 NCCs were replated on ECM deposited by hPSC-ctr-H9 SNs. The number of iPSC-FD-S3 SNs increased as measured by immunofluorescence and flow cytometry (**Fig. 4E and F**). As controls, we also isolated ECM deposited by NCCs and SNs differentiated from iPSCs of FD patients with mild (iPSC-FD-M2) and severe (iPSC-FD-S3) symptoms. We found that ECM from iPSC-FD-M2, but not iPSC-FD-S3, rescued the phenotypes similar to hPSC-ctr-H9 ECM (**Fig. 4C-F**). These results could be explained by the fact that both iPSC-FD-M2 and hPSC-ctr-H9 cells express WT *LAMB4*, whereas iPSC-FD-S3 cells have a variant in *LAMB4*^36^. Therefore, we demonstrate that a healthy ECM is critical for NCC and SN development.

**Figure 4.**
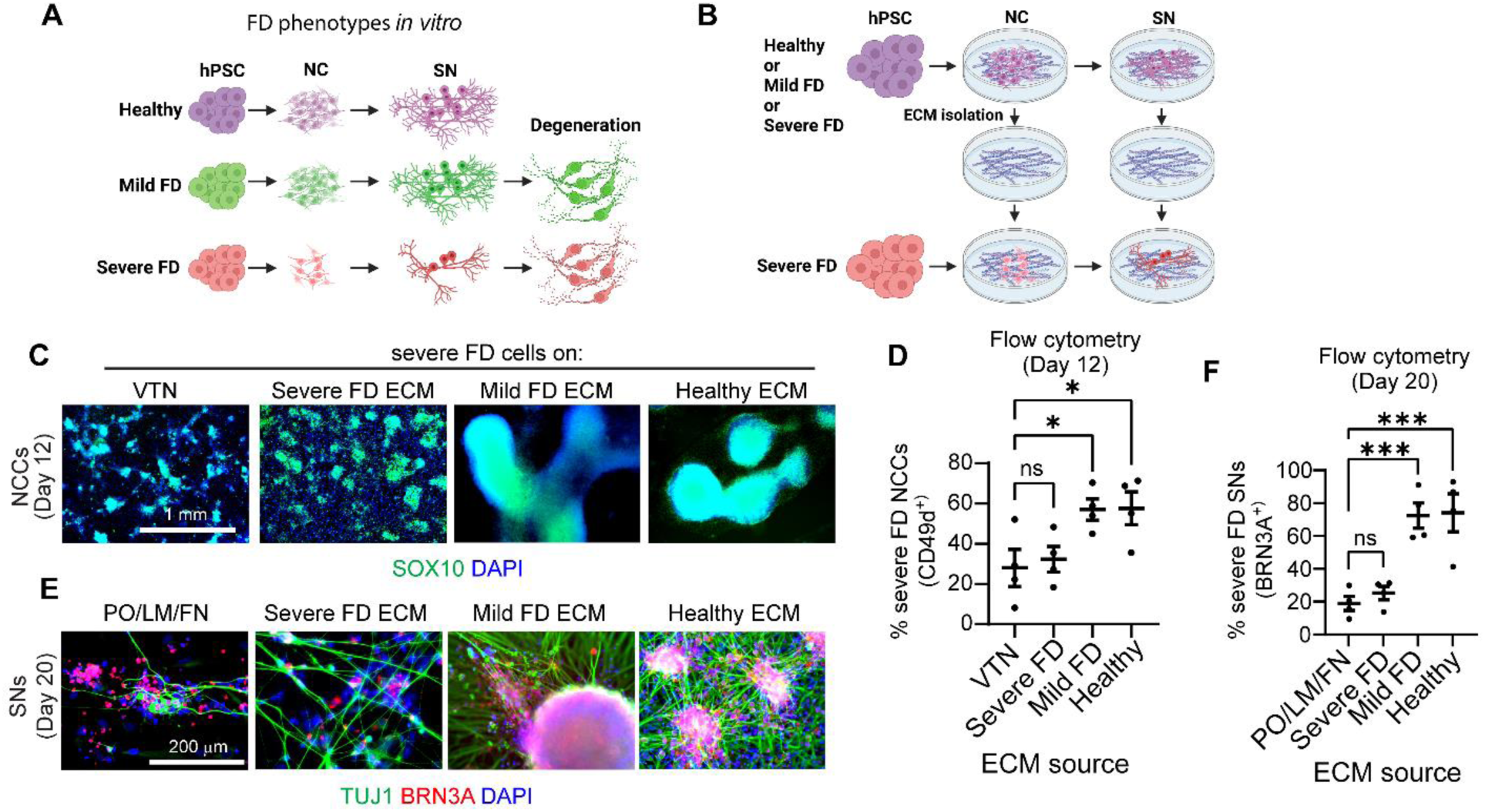
Healthy ECM rescues the developmental defects of SNs from severe FD iPSCs. **A)** Phenotypes observed in SNs differentiated from iPSCs from FD patients with mild and severe FD symptoms^36^. **B)** Schematic of the experimental approach. **C)** Effects of the ECM in NCC differentiation. iPSC-FD-S3 iPSCs were differentiated on vitronectin (VTN) alone or on the isolated ECM deposited by healthy, mild FD, or severe FD NCCs. NCCs were fixed on day 12 and stained for SOX10 and DAPI. **D)** Quantification of NCCs from **C)**. NCCs were harvested and stained for the NCC marker CD49d and analyzed using flow cytometry (n=4 biological replicates). **E)** Effects of the ECM on SN differentiation. iPSC-FD-S3 hPSCs were differentiated in dishes coated with Poly-L-Ornithine (PO), laminin-111 (LM), and fibronectin (FN) or on isolated ECM from healthy, mild FD, or severe FD SNs. SNs were fixed on day 20 and stained for the neuronal marker TUJ1 and SN marker BRN3A. **F)** Quantification of SNs from **E)**. SNs were resuspended and fixed on day 20. Followed by staining for BRN3A and analysis using flow cytometry (n=4 biological replicates). For **D)** and **F),** one-way ANOVA followed by Dunnett’s multiple comparisons test. ns, non-significant, *p<0.05, ****p<0.0001. Graphs show mean ± SEM.

### *LAMB4* is downregulated in sensory neurons affected by severe Familial Dysautonomia symptoms

Our results suggest that the ECM, particularly *LAMB4*, plays a very important role in the development of SNs. Since two *LAMB4* single nucleotide variants have been identified in patients with severe FD symptoms^36^ (**Fig. 5A**), we decided to investigate what are the consequences of these variants in *LAMB4* expression. We used three iPSC lines from severe FD patients previously characterized^36^: iPSC-FD-S1, iPSC-FD-S2, and iPSC-FD-S3. As controls we used a healthy iPSC line (iPSC-ctr-C1) and an iPSC line derived from a patient with mild FD symptoms (iPSC-FD-M2), both of which do not harbor any variants in *LAMB4*. We first measured *LAMB4* expression during development. Similarly to the control hPSC-ctr-H9 cells, iPSC-ctr-C1 and iPSC-FD-M2 cells expressed *LAMB4* starting at the late stages of NCC differentiation and the early stages of SN specification (**Fig. 5B**). However, *LAMB4* expression peaked on day 16, instead of day 20 in hPSC-ctr-H9 (**Fig. 1D**), possibly due to intrinsic differences between iPSCs and human embryonic stem cells (hPSC-ctr-H9). Interestingly, the three severe FD lines showed lower *LAMB4* expression compared to the controls (**Fig. 5B and C**). We next tested whether transcriptional downregulation was also reflected at the protein level. We measured the expression of laminin β4 from cell lysates from SNs from day 20 to day 50 of the differentiation and found that it followed the same pattern as hPSC-ctr-H9 SNs. In the control line, low levels of laminin β4 were detected on day 20, which then increased until day 50, possibly due to the deposition of laminin β4 in the ECM (**Fig. 5D and E**). In contrast, laminin β4 levels did not increase in the severe FD SNs, suggesting that the variants observed in these lines affect *LAMB4* transcription and subsequent translation (**Fig. 5D and E**). We confirmed these observations by immunofluorescence (**Fig. 5F and G**), where the signal intensity of laminin β4 in iPSC-ctr-C1 and iPSC-FD-M2 SNs was higher compared to iPSC-FD-S2 SNs (**Fig. 5H**). Finally, we measured the levels of laminin β4 in the ECM and confirmed that iPSC-FD-S2 SNs expressed lower laminin β4 levels compared to iPSC-ctr-C1 SNs (**Fig. 5I and J**).

**Figure 5.**
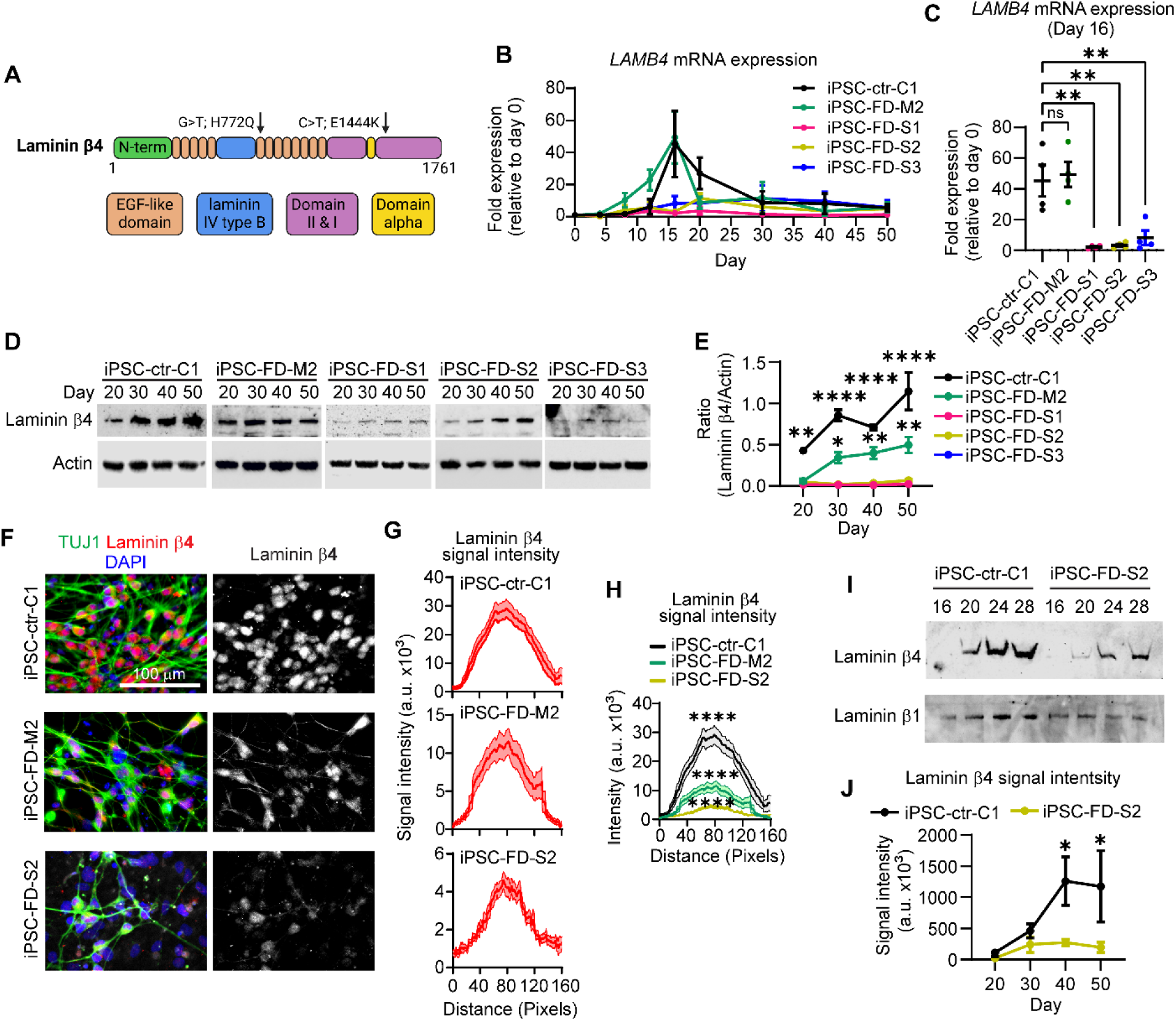
*LAMB4* expression is downregulated in patients with severe FD symptoms. **A)** Schematic of the single nucleotide variants identified in *LAMB4* in patients with severe FD^36^. **B)** *LAMB4* expression in SNs differentiated from iPSCs of patients with severe FD. Severe FD iPSC lines S1, S2, and S3, one mild FD iPSC line (M2), and one healthy control iPSC line (C1) were differentiated into SNs. Total RNA was isolated in the indicated times and gene expression was measured by RT-qPCR (n=4 biological replicates). **C)** *LAMB4* expression by SNs on day 16 shown in **B)** is shown (n=4 biological replicates). **D)** Laminin β4 expression during SN development. iPSC-FD-S1, iPSC-FD-S2, iPSC-FD-S3, iPSC-FD-M2, and iPSC-ctr-C1 cells were differentiated into SNs. Lysates were collected on the indicated days and immunoblotted for laminin β4 and actin. **E)** Quantification of signal intensity of immunoblots shown in **D)** (n=3 biological replicates). Difference between iPSC-FD-S1, iPSC-FD-S2, and iPSC-FD-S3 vs iPSC-ctr-C1 or iPSC-FD-M2 was analyzed. **F)** Laminin β4 in SNs. iPSC-FD-S2, iPSC-FD-M2, and iPSC-ctr-C1 cells were differentiated into SNs. Cells were fixed on day 20 and stained for laminin β4, TUJ1, and DAPI. **G)** Laminin β4 signal intensity measured from images in **F)**. The average of 20 cells is plotted (n=3 biological replicates). **H)** Comparison of laminin β4 signal intensity from **G)**. **I)** Laminin β4 levels in the ECM of severe FD SNs. ECM deposited by SNs from iPSC-FD-S2 and iPSC-ctr-C1 cells was isolated on the indicated days and immunoblotted for laminin β4. Plates were coated with laminin β1 which was used as a loading control. **J)** Signal intensity of blots from **I)** (n=4 biological replicates). For **C)**, one-way ANOVA followed by Tukey’s multiple comparisons test. For **E)** and **J)**, two-way ANOVA followed by Šídák’s multiple comparisons test. For **H)**, two-way ANOVA followed by Tukey’s multiple comparisons tests. ns, non-significant, *p<0.05, **p<0.005, ****p<0.0001. Graphs show mean ± SEM.

We next asked whether restoring *ELP1* expression rescues the expression of *LAMB4* in FD. To answer this question we used two previously characterized FD iPSC lines (iPSC-rescued-T6.1 and iPSC-rescued-T6.5) where the *ELP1* mutation was rescued^36^. However, since they were generated from iPSC-FD-S2 cells, they still harbor the *LAMB4* variant identified in this cell line^36^. Similar to the parental line (iPSC-FD-S2), iPSC-rescued-T6.1 and iPSC-rescued-T6.5 cells expressed lower levels of *LAMB4* mRNA compared to iPSC-ctr-C1 cells (**Fig. S5A and B**). This was confirmed by immunoblotting of the total levels of laminin β4 as well as immunofluorescence (**Fig. S5C-E**). Together, our results suggest that decreased *LAMB4* expression in addition to the *ELP1* mutation in FD may cause severe symptoms, and therefore has clinical implications. Moreover, *LAMB4* could be used as a marker to detect early onset of severe symptomatology in FD and thus be used in personalized medicine.

### Laminin β4 forms laminin-443 and controls actin filament accumulation in sensory neurons

Since *LAMB4*/laminin β4 is necessary for SN development and has clinical implications, we decided to gain knowledge into how laminin β4 regulates SN development. Laminins are assembled in trimers consisting of chains α, β, and γ^15^ (**Fig. 6A**). There are five known α chains, four β chains, and three γ chains. To date, no laminin trimer containing laminin β4 has been described, thus, we first asked which chains interact with it. We measured the expression of every laminin chain in day 16 SNs and found that in addition to *LAMB4*, *LAMA4* and *LAMC3* were upregulated (**Fig. 6B**). *LAMA4* encodes laminin α4 and *LAMC3*, laminin γ3, which suggests that laminin β4 is part of the laminin-443. To confirm this, we immunoprecipitated laminin α4 and found that laminin β4 and laminin γ3 came down as a complex (**Fig. 6C and D**). Next, we investigated the communication of the laminin β4-containing laminin trimer into the cell during SN development. Laminins bind to integrins at the plasma membrane, which through interactions with talin and vinculin, ultimately control the actin cytoskeleton (**Fig. 6E**). Through this mechanism, laminins ultimately regulate many cellular processes such as cell migration^3^. Based on the literature and our results showing that loss of *LAMB4* affects NCC migration, we hypothesized that laminin β4 also controls actin in SNs. We looked at the expression of F-actin using confocal microscopy in SNs differentiated from *LAMB4^+/+^*, *LAMB4^+/−^*, and *LAMB4^−/−^* hPSCs. F-actin signal was evenly distributed around the cell body of *LAMB4^+/+^* SNs, however the expression changed into a slight punctuated pattern in *LAMB4^+/−^* SNs, and this pattern became more prominent in *LAMB4^−/−^* SNs (**Fig. 6F**). This change also correlated with a decrease in F-actin signal intensity (**Fig. 6G**), suggesting that laminin β4 regulates the formation of F-actin in SNs. Vinculin localization also changed upon loss of *LAMB4* (**Fig. 6H and I**). Vinculin was detected throughout the cytoplasm of *LAMB4^+/+^* SNs, possibly due to its localization at focal adhesions mediating the interaction between the cells and the ECM deposited on the surface of the well. In contrast, *LAMB4^+/−^* SNs showed vinculin accumulation at the plasma membrane in addition to the cytoplasm. Finally, cytoplasmic vinculin localization was lost in *LAMB4^−/−^* SNs and it was present mainly at the plasma membrane (**Fig. 6H and I**). These results show that the laminin β4, via laminin-443, maintains expression of F-actin at the cell body of SNs and transduces biophysical cues to the cell via vinculin. Moreover, in the absence of laminin β4, vinculin is no longer localized in the cytoplasm, potentially dissociated from focal adhesions and F-actin, thus affecting cell migration and differentiation. Our studies highlight a direct link between the cellular environment and intracellular components, and the critical role it plays during development which could potentially be targeted to treat peripheral neuropathies.

**Figure 6.**
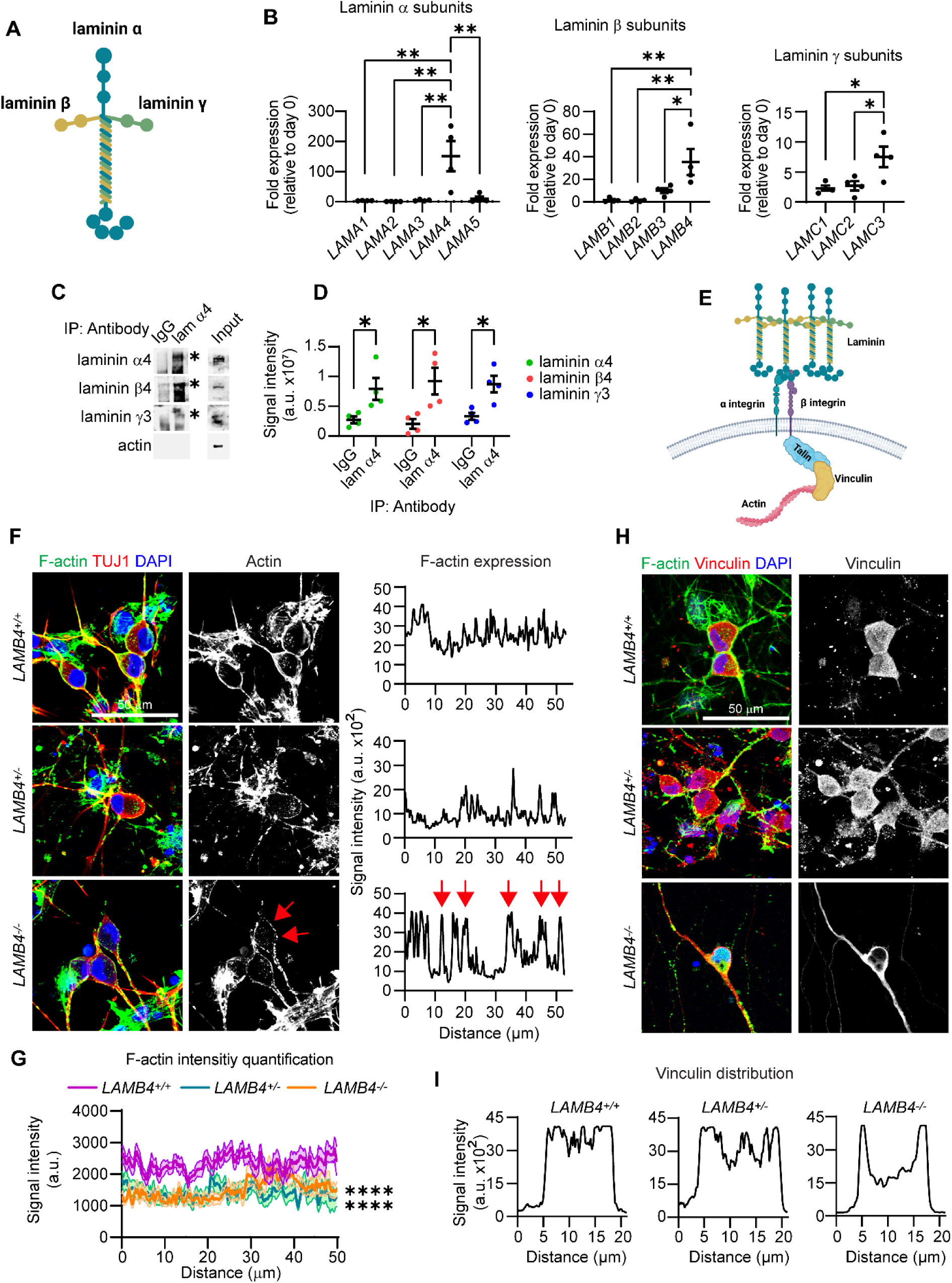
Laminin β4 interacts with laminin α4 and laminin γ3 and regulates actin filament (F-actin) expression. **A)** Schematic of laminin trimer. **B)** Expression of laminin chains. RNA of day 16 hPSC-ctr-H9 SNs was isolated, and mRNA expression of all laminin chains was assessed by RT-qPCR (n=4 biological replicates). **C)** Immunoprecipitation of laminin α4. Lysates from hPSC-ctr-H9 SNs was collected on day 30, followed by immunoprecipitation of laminin α4 (lam α4) and immunoblotting for laminin α4, laminin β4, and laminin γ3. Asterisks mark the band corresponding for each protein. **D)** Quantification of signal intensity of immunoblots in **C)** (n=4 biological replicates). **E)** Schematic of regulation of intracellular pathways by laminins. **F)** Effects of *LAMB4* downregulation on F-actin. *LAMB4^+/+^*, *LAMB4^+/−^*, and *LAMB4^−/−^* SNs were fixed on day 20 and stained for F-actin (Phalloidin), TUJ1, and DAPI (left). Signal intensity of F-actin around the cell body from a representative experiment was measured and plotted (right). Red arrows indicate signal from actin puncta. **G)** F-Actin signal intensity of images from 20 cells in **F)** (n=3 biological replicates). **H)** Vinculin localization upon of loss *LAMB4*. *LAMB4^+/+^*, *LAMB4^+/−^*, and *LAMB4^−/−^* SNs were fixed on day 20 and stained for F-actin (Phalloidin), Vinculin, and DAPI. **I)** Signal intensity of vinculin from a representative experiment from **H)** was measured and plotted. For **B)**, one-way ANOVA followed by Tukey’s multiple comparisons test. For **D)**, two-tailed t-test. For **G)**, two-way ANOVA followed by Tukey’s multiple comparisons test. *p<0.05, **p<0.005, ****p<0.0001. Graphs show mean ± SEM.

## Discussion

*LAMB4*/Laminin β4 has been vastly understudied. We found *LAMB4* orthologs in many species, including humans, chicken, zebrafish, and frogs, but not in rodents (**Fig. S1A**). In addition, we show that *LAMB4* is expressed only in ectoderm lineages derived from NCCs and for a short period of time: 1) during the late stages of migratory NCC and 2) in early stages of SN specification (**Fig. 1D**). These factors make *LAMB4* a difficult gene to study, as it cannot be studied in mouse models, and it is not widely expressed.

*LAMB4* expression in late-stage NCCs and early-stage SNs suggests that it is important for the development of NCC-derived tissues including SNs. A report showing that laminin β4 is expressed in the cutaneous basement membrane^38^ and our results showing that *LAMB4* downregulation reduces NCC migration (**Fig. 2E-H**) support this hypothesis. This timing also explains our gene expression results, where *SOX10* expression in NCCs does not change in the absence of *LAMB4*. In contrast, genes expressed after *SOX10* such as *P75NTR*, *NGN1*, and *NGN2*, which induce NCCs into SN lineages^6,28^, are downregulated. This is not unexpected, as other laminin chains also regulate NCC migration *in vivo*^39^. In SNs, accumulation of laminin β4 in the ECM is visible over 20 days after mRNA is downregulated. It is possible that the spike of *LAMB4* transcription causes a burst of laminin β4 translation and secretion, which is necessary to further differentiate NCCs into SNs. The ECM is required for multiple aspects of development, maturation, and function of neurons^30^, including neurotransmission^40^, the activity of neuromodulators^37^, and promotes synaptogenesis^41^. Moreover, the ECM provides the necessary cues required for axon elongation and guidance during development and after injury^30,42,43^. We observed that axons of *LAMB4^−/−^* SNs show an irregular elongation pattern compared to control SNs (**Fig. 3B**), suggesting that laminin β4 is necessary for this process. These deficiencies in axon elongation could also affect SN homeostasis and explain why *LAMB4^−/−^* SNs degenerate faster than healthy SNs (**Fig. 3I**). This also suggests that laminin β4 secreted by differentiated SNs is required for their survival and agrees with the literature showing that neurons release laminins^41,44–46^.

*LAMB4* downregulation is linked to diseases of the peripheral nervous system (peripheral neuropathies). Patients with sporadic cases of the enteric peripheral neuropathy diverticulitis have been shown to also harbor *LAMB4* variants resulting in its downregulation^16^. Diverticulitis is caused by reduced neuronal density of NCC-derived enteric neurons^47,48^. *LAMB4* has also been linked to the peripheral neuropathy FD. It was previously reported that patients with severe, but not mild FD symptoms harbor mutations in *LAMB4*^36^. We show that SNs differentiated from iPSCs reprogramed from patients with severe FD had lower *LAMB4* expression (mRNA and protein) compared to mild FD SNs (**Fig. 5D-H**). Approximately 99.5% of patients with FD have a mutation in *ELP1*^32^, *LAMB4* expression could explain the symptomatic differences between patients with mild and severe symptoms. Moreover, our results hint at the possibility that *LAMB4* could be used as a diagnostic marker for severe FD onset early in life.

Restoring *ELP1* expression in severe FD iPSCs did not impact *LAMB4* expression, suggesting that *LAMB4* expression is independent of *ELP1*. ELP1 is the scaffold protein of the elongator complex, and it is involved in transcription and tRNA modification during translation^49^. Our results showing that mild, but not severe FD SNs express high levels of *LAMB4* further hints at the possibility that *LAMB4* expression is not completely *ELP1*-dependent. Another possibility is that ELP1 primarily impacts laminin β4 translation due to defects in tRNA production. Thus, laminin β4 translation could be downregulated in mild FD SNs due to the ELP1 mutation, whereas in severe FD SNs, *LAMB4* mRNA expression (due to the identified *LAMB4* mutations) and translation (due to the *ELP1* mutation) are both affected. Further studies will be necessary to dissect this mechanism.

Our studies show that laminin β4 is part of laminin-443. Laminin α4 has been shown to be expressed in the dorsal root ganglia (where NCCs further develop into SNs) during mouse development^50^ which strengthens the hypothesis that laminin β4 is involved in SN development. However, because *LAMB4* is not expressed in rodents, it would be necessary to confirm this possibility in other *in vivo* models. Laminins bind to integrins located on the cell surface and connect to the actin cytoskeleton^51^. Laminin α4 has been shown to interact with integrin α3β1 and α6β1^52,53^, however this interaction has not been explored in SNs. Our results suggest that laminin β4 activates integrins and cause the formation of F-actin around the cell body of SNs. Loss of *LAMB4* results in reduced F-actin expression, possibly due to integrin inactivation. Interestingly, vinculin localization is also altered in *LAMB4^−/−^* SNs. Vinculin is a cytoplasmic protein that links integrins to actin and translates biophysical cues from the ECM into intracellular biochemical signals^54^. Upon integrin activation, vinculin is recruited to F-actin and focal adhesions^51,54,55^. Loss of *LAMB4* resulted in changes in vinculin localization, possibly due to the inactivation of integrins. In this scenario, it is possible that vinculin associates with other known interactors present at the cell membrane, such as phosphatidylinositol 4,5-bisphosphate^56^. These are interesting observations, as most of the studies of actin in neurons focus on the growth cone^57^ and opens new avenues to study the cell biology of peripheral neurons. Furthermore, understanding ECM and actin regulation in the peripheral nervous system will uncover new mechanisms that could be used to promote neuronal regeneration and treat peripheral neuropathies.

## Materials and methods

### hPSC maintenance

hPSC-ctr-H9 human embryonic stem cells (WA-09, WiCell) and all human induced pluripotent stem cells were grown at 37 °C with 5 % CO_2_ in vitronectin-coated dishes (ThermoFisher, cat# A31804, 5 μg/mL, 1 h at RT). Cells were fed daily with Essential 8 Medium + Supplement (Gibco, cat# A1517001). Cells were split at a 1:10 ratio using the following protocol: cells were washed with PBS, incubated with 0.5 mM EDTA, 3.08 M NaCl in PBS with for 2 minutes at 37 °C, and then resuspended in E8 + Supplement. iPSC-ctr-C1, iPSC-FD-M2, iPSC-FD-S1, iPSC-FD-S2, and iPSC-FD-S3 were previously characterized^36^.

### Sensory neuron differentiation

Differentiation was done as previously described^23,24^. Prior to differentiation, plates were coated with vitronectin (5 μg/mL) and incubated for 1h at RT. On day of plating (day 0), hPSCs were washed with PBS, incubated with 0.5 mM EDTA, 3.08 M NaCl in PBS for 20 minutes, and plated at a density of 200,000 cells/cm^2^ in NC differentiation media (day 0-1) containing: Essential 6 Medium (Gibco, cat# A1516401), 10 μM SB431542 (R&D Systems, cat# 1614), 1 ng/mL BMP4 (R&D Systems, cat# 314-BP), 300 nM CHIR99021 (R&D Systems, cat# 4423), and 10 μM Y-27632 (Biogems, cat# 1293823). BMP4 concentration was titrated for each line. Accordingly, BMP4 was not used with iPSC-FD-S1 and iPSC-FD-S3 cells. The next day, the cells were fed with NC differentiation media (day 0-1). From day 2 to 12, cells were fed every two days with NC differentiation media (day 2-12) containing: Essential 6 Medium, 10 μM SB431542, 0.75 μM CHIR99021, 2.5 μM SU-5402 (Biogems, cat# 2159233), and 2.5 μM DAPT (R&D Systems, cat# 2634).

On day 10, plates were coated with 15 μg/ml poly-L-ornithine (PO, Sigma, cat# P3655) in PBS and incubated at 37 °C overnight. On day 11, the plates were washed 3X with PBS and coated with 2 μg/ml laminin-1 (LM, Cultrex, cat# 3401-010-02) and 2 μg/ml human fibronectin (FN, Corning, cat# 47743-654) in PBS and incubated overnight. On day 12, cells were resuspended using Accutase (Innovative Cell Technologies, cat# NC9464543) for 20 minutes, washed with PBS, and resuspended in SN Media containing Neurobasal media (Gibco, cat# 21103-049) containing 1X N2 (Gibco, cat# 17502-048), 1X B-27 (Gibco, cat# 12587-010), 2 mM L-glutamine (ThermoFisher, cat# 25030-081), 20 ng/ml GDNF (Peprotech, cat# 450-10), 20 ng/ml BDNF (R&D Systems, cat# 248-BD), 25 ng/ml NGF (Peprotech, cat# 450-01), 600 ng/ml of laminin-1, 600 ng/ml fibronectin, 1 μM DAPT and 0.125 μM retinoic acid (Sigma, cat# R2625). Cells were then replated at a density of 250,000 cells/cm^2^ onto PO/LM/FN coated plates. The media was replaced the following day. Cells were fed every 2-3 days. On day 20, DAPT was removed. Differentiation progress was followed using a brightfield microscope (Leica).

### Endoderm differentiation

Endoderm differentiation was performed as described^36,5831,44^. On day 0, hPSC-ctr-H9 cells were washed with PBS and incubated with Accutase for 20 min and seeded at a density of 100,000 cells/cm^2^ in RPMI medium (ThermoFisher, cat# 12633012) with Glutamax (ThermoFisher, cat# 35050061) and 100 ng/mL Activin A (R&D Systems, cat# 338-AC-010). Cells were fed daily for 3 days and FBS was added at increasingly concentrations: 0%, 0.2%, and 2%.

### Mesoderm and cardiomyocyte differentiation

Cardiomyocyte differentiation was done as previously described^59^. hPSC-ctr-H9 colonies were washed with PBS followed by incubation with Accutase for 20 min. Cells were resuspended in E8 medium + supplement and seeded at a density of 100,000 cells/cm^2^. When the cells reached ∼80% confluency, the cells were fed with RPMI medium supplemented with insulin-free B27 (ThermoFisher, cat# A1895601) and 6 µM CHIR99021 for 2 days. A day later, the media was replaced with RPMI + insulin-free B27. On day 4, cells were fed with RPMI + insulin-free B27 with 5 µM IWP2 (Cayman Chemical, cat# 13951). The following day, the media was replaced with RPMI + insulin-free B27. The cells were fed on day 7 with RPMI + insulin-free B27 and media was replaced every 2 days.

### RNA isolation and RT-qPCR

RNA was isolated using Trizol (ThermoFisher, cat# 15596026) according to the manufacturer’s conditions and resuspended in 20 μL RNase-free water. RNA concentration and purity was measured using NanoDrop One (ThermoFisher). 1 μg of RNA was converted to cDNA using iScript cDNA Synthesis kit (BioRad, cat# 1708841) according to the manufacturer’s instructions and diluted 1:100 in RNase-free water. RT-qPCR reactions were run with 1 ul of cDNA and SYBR Green Supermix (BioRad, cat# 1725272) according to the manufacturer’s conditions in a C1000 Touch Thermal Cycler CFX96 (BioRad). The following cycling parameters were used: 95°C for 5 minutes, 40 cycles of 95°C for 5s and 60°C for 10 s. Results were analyzed using the comparative CT method. GAPDH was used as a housekeeping gene. The sequences of primers used in this study are available in Supplementary Table 1.

### Antibodies

Laminin β4 (Abcam, cat# ab150819; Sigma, cat# HPA020242), laminin β1 (Abcam, cat# ab44941), laminin α4 (R&D Systems, cat# AF7340), laminin γ3 (Proteintech, cat# 67261-1-I), SOX10 (Santa Cruz, cat# sc-365692), TFAP2A (Abcam, cat# ab108311), BRN3A (Millipore, cat# MAB1585), TUJ1 (Biolegend, cat# 801201), ISL1 (DSHB, cat# 39.4D5-c), PRPH (Santa Cruz, cat# sc-377093), Actin (BD Biosciences, cat# 612656), Vinculin (Abclonal, cat# A14193), αSMA (Sigma, cat# A5228), Phalloidin-iFluor 488 (Abcam, cat# ab176753), CD49d-PE/Cy7 (Biolegend, cat# 304314), TRKA-PE (R&D Systems, cat# FAB1751P), TRKB-AF647 (R&D Systems, cat# FAB3971R), and TRKC-PE (R&D Systems, cat# FAB373P). The following secondary antibodies were used: From ThermoFisher: goat anti-mouse IgG1 AF488 (cat# A21121), goat anti-mouse IgG2a (cat# A-21131), goat anti-mouse IgG2b (cat# A21242), donkey anti-rabbit AF647 (cat# A31573), donkey anti-mouse AF488 (cat# A21202), goat anti-mouse HRP (cat# 62-6520), and goat anti-rabbit HRP (cat# 65-6120), Goat anti-rat HRP (cat# A18865). Donkey anti-sheep HRP antibody (Jackson Immunoresearch, cat# 713-035-003). The dilutions used are indicated in each section.

### Immunoblotting

To collect cell lysates, cells differentiated in 6-well plates were washed with PBS and incubated with 120 µL of RIPA buffer (Sigma, cat# R0278) with 1 mM PMSF and 1X PhosSTOP (Roche, cat# 4906845001) for 15 minutes on ice. Cells were then scrapped and the lysate transferred to an Eppendorf tube, followed by mixing 10 s using a vortex and centrifuged at 12,000 RPM for 10 minutes at 4°C. Supernatants were transferred to a new Eppendorf tube and protein concentration was measured. Samples were mixed with 2X Laemmli buffer containing β-mercaptoethanol and ran in 7.5% polyacrylamide gels under denaturing conditions using MOPS buffer at 130 V. Proteins were transferred to a nitrocellulose membrane and blocked for 30 minutes in 5% non-fat dry milk in 0.1 % TBS-T (0.1% Tween-20, 50 mM Tris-HCl, 150 mM NaCl, pH7.6). Primary antibodies were added to the membranes in blocking buffer (laminin β4 – 1:1000, laminin α4 – 1:1000, laminin γ4 – 1:1000, Actin – 1:5000) and incubated overnight at 4 °C. Blots were then washed 3X with 0.1 % TBS-T and incubated with goat anti-mouse HRP, goat anti-rabbit HRP, goat anti-rat HRP, or donkey anti-sheep HRP antibody (1:5000) for 1 h at room temperature. Blots were washed 3X with 0.1% TBS-T and incubated with Clarity Western ECL Substrate (BioRad, cat# 1705061). Chemiluminescence signal was detected using UVP ChemStudio (Analytic Jena). Signal quantification was done using Image Studio Lite (LICOR).

### Immunoprecipitation

Lysates were collected and concentration was measured as described above. Magnetic protein A/G beads (25 µL, ThermoFisher, cat# 88802) were pre-washed 3X with RIPA buffer with 1 mM PMSF and 1X PhosphoSTOP and incubated with 1 µg of laminin α4 antibody for 30 minutes at 4 °C in a rotator. Beads were then washed 3X with RIPA buffer with 1 mM PMSF and 1X PhosphoSTOP and incubated overnight with 1 mg of lysate. The following day, beads were washed 3X with RIPA buffer with 1 mM PMSF and 1X PhosphoSTOP, and resuspended in 2X Laemmli buffer.

### Immunofluorescence

NCCs and SNs differentiated in 24-or 4-well plates were washed once with PBS and fixed with 4% paraformaldehyde (ThermoFisher, cat# AAJ19943K2) for 20 minutes at RT. Cells were then washed with PBS and incubated for 20 minutes with Permeabilization buffer containing 1% BSA, 0.3% Triton-X, 3% goat or donkey serum and 0.01% sodium azide in PBS. Cells were then incubated with the indicated primary antibodies (laminin β4 – 1:100, SOX10 – 1:100, TFAP2A – 1:500, BRN3A – 1:100, TUJ1 – 1:1500, ISL1 – 1:200, PRPH – 1:100, αSMA – 1:100) in Antibody buffer containing 1% BSA, 3% goat or donkey serum and 0.01% sodium azide overnight at 4°C.

Cells were then washed 3X in PBS and incubated with secondary antibodies in Antibody buffer for 1 h. Cells were washed with PBS, incubated with DAPI (1:1,000) for 5 minutes, washed with PBS, and stored at 4°C. Imaging was done using a Lionheart FX fluorescence microscope (BioTek). Image analyses and quantifications were done in Fiji. For quantifications, 5 different fields were imaged and quantified. For confocal microscopy, 50,000 NCCs were seeded in PO/LM/FN-coated 4-well chamber slides (iBidi, cat# 80426) on day 12. On day 20, SNs were fixed and stained as described above. Primary antibodies used: TUJ1 – 1:1500, Vinculin – 1:100. Phalloidin-iFluor 488 (1:1000) was incubated with secondary antibodies for 1 h. Imaging was done in an Olympus FV1200 Confocal Laser Scanning Microscope using Argon and Helium-Neon lasers. Images were taken as Z-stacks of 3 µm of height. ImageJ was used to obtain maximum intensity projections and to measure the signal intensity profiles.

### Flow cytometry

On the indicated days, cells were washed with PBS and incubated with Accutase for 30 minutes at 37 °C. Cells were then washed and resuspended in Flow buffer (DMEM, 2% FBS, and 1mM L-glutamine) followed by centrifugation at 200 g for 4 minutes. Cells were resuspended in cold PBS, counted, and diluted to a concentration of 1×10^6^ cells/100μL. For NCCs, cells were centrifuged at 200 g for 4 minutes at 4 °C and resuspended in 100 μL of Flow buffer and incubated with CD49d-PE/Cy7 antibody (1:160) for 30 minutes, or with TRKA-PE (1:20), TRKB-AF647 (1:20), or TKC-PE (1:20) antibodies for 1 hour on ice. Samples were washed 2X with Flow buffer, resuspended in 300 μL of Flow buffer with DAPI (1:1000), filtered, and analyzed using a Cytoflex S (Beckman Coulter). For SNs, cells at a concentration of 1×10^6^ cells/100μL were centrifuged, resuspended in 300 μL BD Cytofix buffer (BD Biosciences, cat# 554655), and incubated on ice for 30 minutes. Cells were centrifuged for 4 minutes at 2,000 RPM and resuspended in 600 μL of cold BD Perm/Wash buffer (BD Biosciences, cat# 554723). Goat serum (30 μL) was added to the cells and incubated on ice for 30 minutes. Cells were divided in 3 tubes (200 μL each): 1) unstained control, 2) secondary antibody control, and 3) sample. All tubes were centrifuged for 4 minutes at 2,000 RPM and the cells were resuspended in 200 μL of Antibody buffer (BD Perm/Wash buffer + 10 μL goat serum) with or without BRN3A antibody (1:100) and incubated overnight at 4°C. Cells were then washed twice with 300 μL BD Perm/Wash buffer, resuspended in Antibody buffer with or without AF488 goat-anti-mouse (1:500), and incubated on ice for 30 minutes. Cells were then washed 3X with BD perm/wash buffer, filtered, and analyzed using a Cytoflex S (Beckman Coulter). Analyses were done using FlowJo.

### Scratch assay

On day 8, NCCs differentiated from *LAMB4^+/+^*, *LAMB4^+/−^*, and *LAMB4^−/−^* hPSCs were washed with PBS and incubated with Accutase for 20 minutes at 37 °C. Cells were resuspended in NC differentiation media (day 2-12), counted, and replated at a density of 60,000 cells/cm^2^ in 4-well or 24-well plates. When the cells reached confluency, a scratch was performed in the center of the well using a 1 000 µL sterile tip. Brightfield images were immediately taken (0 h) was taken using a Lionheart FX (Bio-Tek) fluorescent microscope. Subsequent images were taken 24 and 48 h later at the same coordinates. Images were analyzed as previously described^60^.

### Live-cell imaging

On day 8, NCCs from *LAMB4^+/+^*, *LAMB4^+/−^*, and *LAMB4^−/−^* hPSCs were washed with PBS, incubated with Accutase for 20 minutes at 37 °C, and resuspended in NC differentiation media (day 2-12). Cells were then counted and replated at a density of 15,000 cells/cm^2^ in 4-well or 24-well plates. Medium was replaced the following day and brightfield images were taken every 10 minutes for 18 h using a Lionheart FX microscope (Bio-Tek) with climate control chamber. Cells were maintained at 37 °C with 5 % CO_2_ throughout the experiment. Each experiment was performed in triplicate (technical replicate) and approximately 60-80 cells were tracked per well. Individual images were compiled using Fiji and individual cells were tracked using TrackMate (v7.13.2)^61,62^. Tracks of individual cells were exported and analyzed using the Chemotaxis and Migration Tool software (Ibidi).

### Generation of *LAMB4* mutant hPSCs

Two gRNAs (GCTCAAGATGACTGCAACAG and CTGGTGATCTCCTGGTGGGC) targeting exon 3 of *LAMB4* were selected using E-CRISP^63^ (available at www.E-CRISP.org). The oligos were annealed, phosphorylated, and ligated into PX458 using T4 DNA ligase. The resulting plasmid was transformed into DH5α bacteria and colonies were screened by sanger sequencing. The resulting plasmids (PX458-LAMB4gRNA1 and PX458-LAMB4gRNA2) were transfected into hPSC-ctr-H9 cells using Lipofectamine Stem Transfection Reagent (ThermoFisher, cat# STEM00001) following to the manufacturer’s protocol. After 48 h, cells were washed with PBS and incubated with Accutase for 20 minutes at 37 °C. The cells were transferred to a 15 mL conical tube, filled with PBS and centrifuged at 200 g for 5 minutes. The supernantant was aspirated and the pellet was resuspended in sorting medium containing Essential 8 Medium + Supplement, 1X CloneR (Stemcell Technologies, cat# 05889), and 10 μM Y-27632. Cells were then counted and 2×10^6^ cells were transferred to an Eppendorf tube and resuspended in 400 μL of sorting medium containing 0.4 μL of Propidium Iodide (ThermoFisher, Cat# P3566). The resuspended cells were filtered using a round-bottom FACS tube and GFP^+^ cells were sorted using a FACS Melody Cell Sorter System (BD Biosciences). Individual cells were sorted to VTN-coated 96-well plates with prewarmed 50 μL of sorting medium in each well. The cells were fed every 24 h for approximately 10 days. When colonies started to emerge, cells were transferred to 24-well plates using EDTA and the protocol previously described. Genomic DNA was isolated from each clone and screened. Positive clones were further expanded.

### Electrophysiology experiments

Experiments were performed using a Maestro Pro (Axion Biosystems) multi-electrode array (MEA) system. On day 12, NCCs were seeded (250,000 cell/cm^2^) onto PO/LM/FN-coated BioCircuit MEA 96 plates (Axion Biosystems, cat# M768-BIO-96), containing 8 embedded electrodes/well, in SN Media as previously described, and allowed to continue differentiating. Recordings were made every 2-3 days at 37°C with a sampling frequency of 12.5 kHz for 5 minutes. Recordings from at least 6 wells per reading were averaged. Firing frequency was normalized to the number of active electrodes. Bursts were detected using Inter-Spike Interval. Capsaicin (Sigma, cat# M2028) and WIN 55,212-2 (R&D Systems, cat# 1038) were resuspended in DMSO and added to the cells 3 minutes prior to starting recordings. Hypoosmotic media was obtained by mixing SN Media with sterile water in a 45:55 ratio and it was added to the cells prior to recordings.

### Degeneration assay

On day 12, NCCs from *LAMB4^+/+^*, *LAMB4^+/−^*, and *LAMB4^−/−^* hPSCs were replated on 4-well plates (ThermoScientific, cat# 12-565-72), at 250,000 cells/cm^2^, coated with PO/FN in SN media with 1 ng/ml NGF. Cells were fed every 2-3 days. DAPT was removed after day 20. Cells were fixed on day 13, 20, 27, and 34 and stained for BRN3A and TUJ1.

### Extracellular matrix isolation and rescue experiments

NC– and SN-derived ECM was isolated as previously described^25^. To isolate ECM from SNs, day 12, hPSC-ctr-H9, iPSC-ctr-C1, and iPSC-FD-S2 NCCs were resuspended in Accutase as described above and seeded in 60 mm dishes. On day 30, cells were washed with 3 mL of PBS and incubated with 20 mM Ammonium Hydroxide (Sigma, cat# 221228-100ML-A). The dishes were constantly shaken for 5 minutes at RT, followed by 5 washes with 5 mL of de-ionized water. For immunoblotting, the ECM was scrapped and resuspended in Laemmli buffer containing β-mercaptoethanol and 100 mM dithiothreitol (DTT, RPI, cat# D11000) preheated heated at 95 °C for 2 min. For ECM rescue experiments, hPSC-ctr-H9, iPSC-FD-M2, and iPSC-FD-S3 were differentiated using the SN differentiation protocol described above. On day 12 (NCCs) and day 30 (SNs) the cells were treated following the ECM isolation protocol. The undisturbed ECM was kept in the plates in de-ionized water. To start the differentiation, water was aspirated and iPSC-FD-S3 cells were seeded following the SN differentiation protocol described above.

### Bioinformatics

RNAseq data from endoderm^21^ (GSE52658) and mesoderm RNAseq^22^ (GSE85066) were analyzed. FPKM and TPM results were converted to log2 and graphed as heatmaps. For laminin chains analysis the following sequences from NCBI were used: 1) LAMB1: *D. rerio* (NP_775382), *X. tropicalis* (XP_002933140), *M. musculus* (XP_006515056), *R. norvegicus* (XP_003750185), *C. lupus* (XP_038279702), *B. taurus* (NP_001193448), *M. mulatta* (XP_014990159), *H. sapiens* (XP_047276315), *P. troglodytes* (XP_001165667), *G. gallus* (XP_046780211), *A. carolinensis* (XP_016849500), *S. purpuratus* (XP_030828530), *D. melanogaster* (NP_476618), *A. mellifera* (XP_006571829), *C. elegans* (NP_500734); 2) LAMB2: *M. musculus* (NP_001398157), *R. norvegicus* (XP_006243771), *M. mulatta* (XP_014986301), *H. sapiens* (XP_005265184), *P. troglodytes* (XP_016796574), *C. lupus* (XP_038283703), *B. taurus* (XP_010816035), *G. gallus* (NP_989497), *A. carolinensis* (XP_062829843), *D. rerio* (XP_005162102), *X. tropicalis* (XP_004914156), *D. melanogaster* (NP_524006); 3) LAMB3: *D. rerio* (XP_700808.6), *G. gallus* (XP_040547616), *X. tropicalis* (XP_012826649), *A. carolinensis* (XP_062834708), *M. musculus* (XP_006497296), *R. norvegicus (*XP_008768078), *B. taurus* (XP_005217424), *C. lupus* (XP_038526808), *M. mulatta* (XP_014973102), *H. sapiens* (XP_005273181), *P. troglodytes* (XP_054514183); 4) LAMB4: *D. rerio* (XP_068073408), *X. tropicalis* (XP_031754867), *C. lupus* (XP_038310194), *M. mulatta* (XP_028702003), *H. sapiens* (XP_011514277), *P. troglodytes* (XP_063672018), *G. gallus* (XP_040515061), *A. carolinensis* (XP_062837767). Alignments were done using Clustal Omega^64^ using default settings. The phylogenetic tree was visualized using Treeviewer.

### Statistical analysis

All analyses and graphs were done using PRISM (GraphPad). Statistical analyses are indicated in each figure legends. Two-tailed Student’s t-test was used to compare two groups. One-way analysis of variance (ANOVA) followed by Dunnett’s or Tukey’s multiple comparisons test was used to compare three or more groups. Two-way ANOVA followed by Šídák’s multiple comparisons test was used to analyze data sets with two variables. Data presented are shown as mean ± SEM. In all experiments the differences were considered significant when p<0.05. The number of biological replicates (n) are defined as the number of independent differentiations started at least three days apart or from a different vial of cells. The number of biological replicates are indicated in the figure legends.

## Acknowledgments

We thank Dr. Yao Yao (University of South Florida), Dr. Michael Tiemeyer (University of Georgia), and Dr. Natalia Ivanova (University of Georgia) for their input in this project. We thank Dr. Abel Alcazar-Roman (Heinrich Heine University Düsseldorf) for critical reading of the manuscript. We also thank Julie Nelson from the CSRL Cytometry Shared Resource Laboratory (University of Georgia) for her help with flow cytometry experiments. Schematics were done using Biorender.com.

## Author contributions

K.S-D conceived, designed, conducted and analyzed experiments, and wrote the manuscript. T.S., C.J., A.J.P., K.S.T., T.N.K, and S.B.G. conducted experiments. N.Z. conceived, led the study, provided guidance, edited the manuscript, and provided funds.

## Data and material availability

Requests for reagents should be directed to the corresponding author, Nadja Zeltner

