## Supplementary Figures and Tables for "Laminin β4 is required for the development of human peripheral sensory neurons"

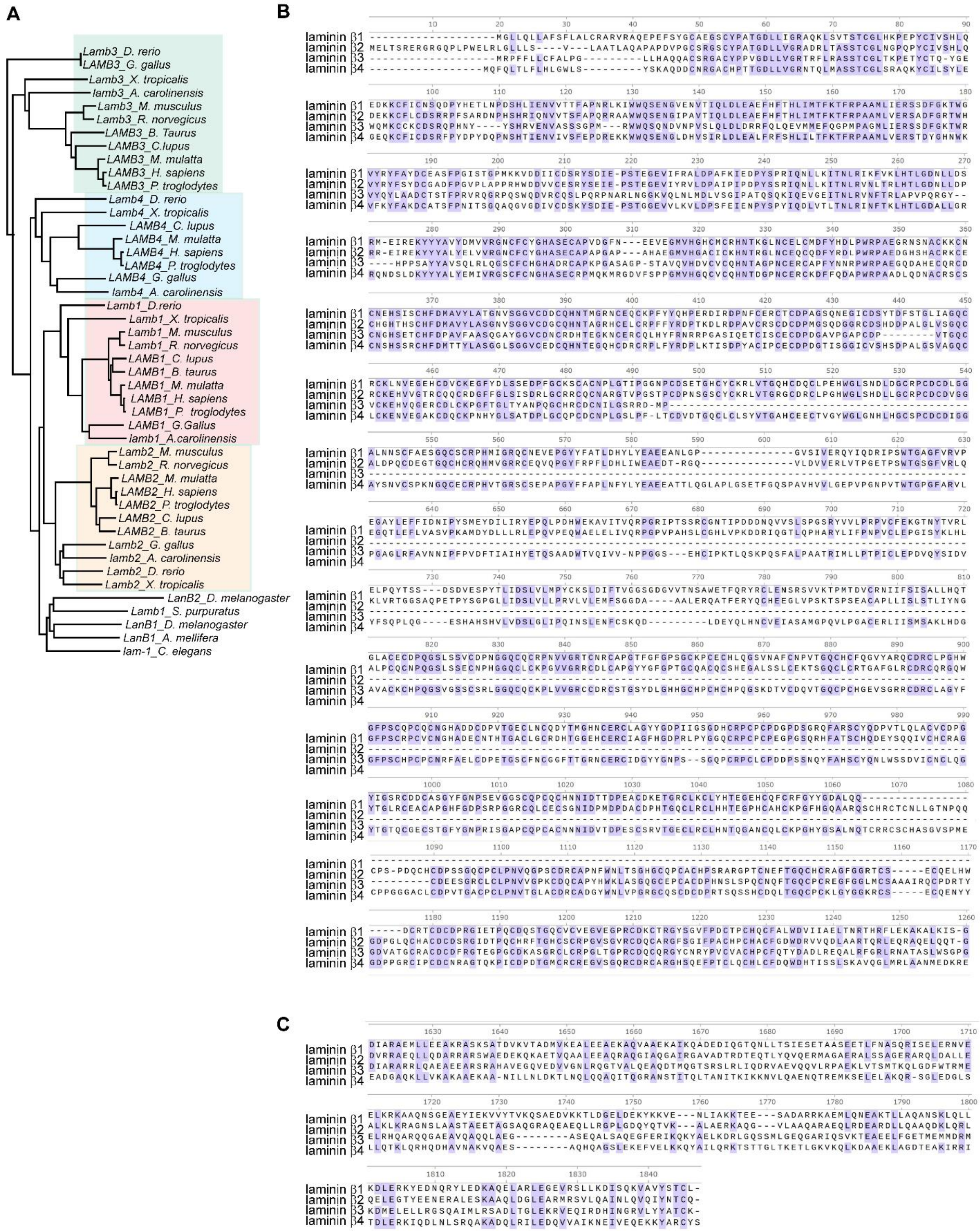

**Figure S1. Similarities between *LAMB* genes. A)** Dendrogram of the *LAMB* genes among species. **B and C)** Alignment of regions in the **B)** N-terminal and **C)** C-terminal domains of the amino acid sequences of the human laminin β chains.

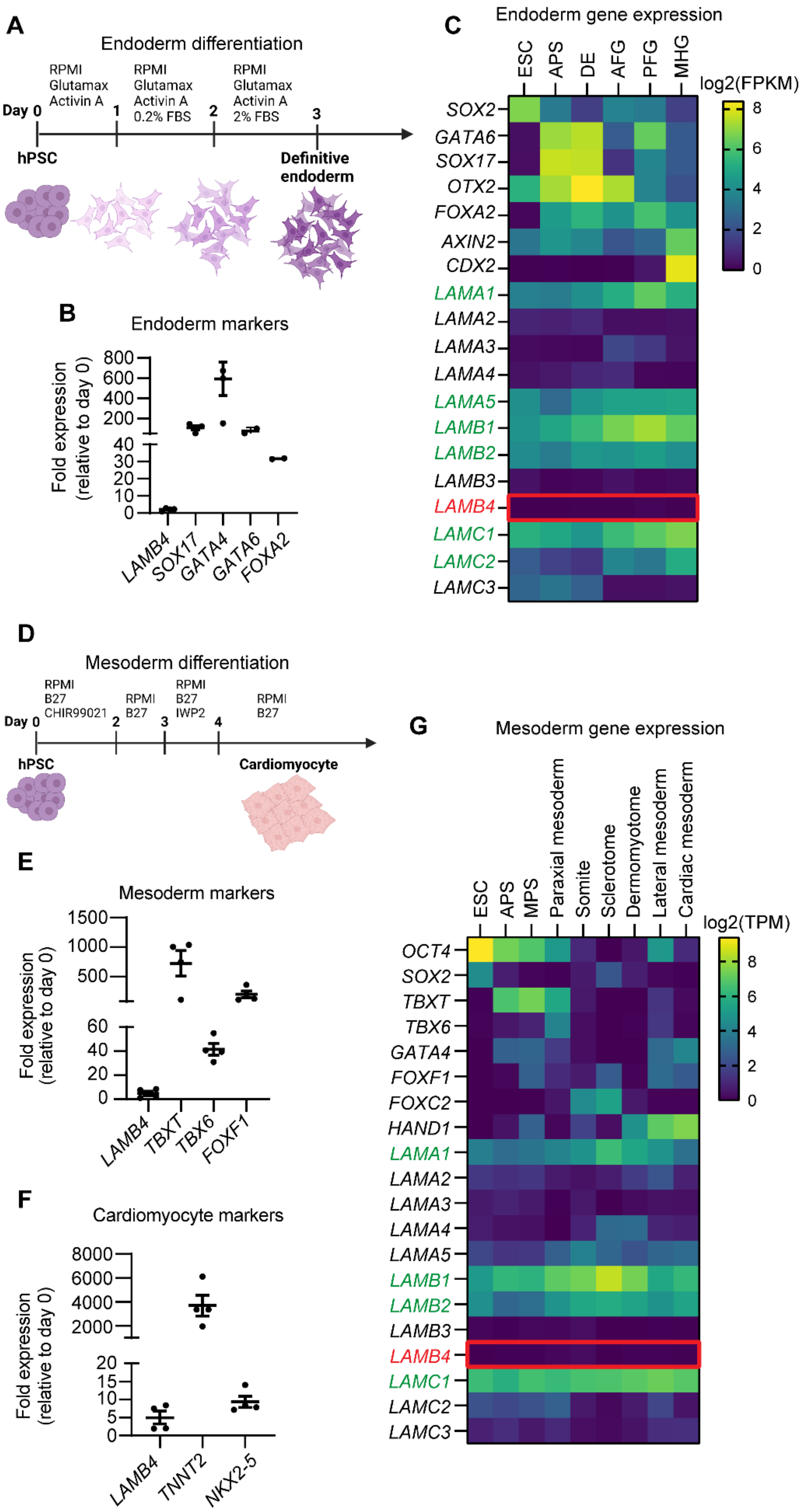
A
Endoderm differentiation
RPMI Glutamax Activin A
RPMI Glutamax Activin A 0.2% FBS
RPMI Glutamax Activin A 2% FBS
Day 0
1
2
3
hPSC
Definitive endoderm
B
Endoderm markers
Fold expression (relative to day 0)
LAMB4
SOX17
GATA4
GATA6
FOXA2
C
Endoderm gene expression
ESC
APS
DE
AFG
PFG
MHG
log2(FPKM)
SOX2
GATA6
SOX17
OTX2
FOXA2
AXIN2
CDX2
LAMA1
LAMA2
LAMA3
LAMA4
LAMA5
LAMB1
LAMB2
LAMB3
LAMB4
LAMC1
LAMC2
LAMC3
D
Mesoderm differentiation
RPMI B27 CHIR99021
RPMI B27
RPMI B27 IWP2
RPMI B27
Day 0
2
3
4
hPSC
Cardiomyocyte
E
Mesoderm markers
Fold expression (relative to day 0)
LAMB4
TBXT
TBX6
FOXF1
F
Cardiomyocyte markers
Fold expression (relative to day 0)
LAMB4
TNNT2
NKX2-5
G
Mesoderm gene expression
ESC
APS
MPS
Paraxial mesoderm
Somite
Sclerotome
Dermomyotome
Lateral mesoderm
Cardiac mesoderm
log2(TPM)
OCT4
SOX2
TBXT
TBX6
GATA4
FOXF1
FOXC2
HAND1
LAMA1
LAMA2
LAMA3
LAMA4
LAMA5
LAMB1
LAMB2
LAMB3
LAMB4
LAMC1
LAMC2
LAMC3

6

**Figure S2. *LAMB4* is not expressed in mesoderm and endoderm. A)** Schematic of the endoderm differentiation protocol. **B)** Gene expression of endoderm markers. hPSC-ctr-H9 cells were differentiated into endoderm and RNA was isolated on day 3. mRNA levels were measured using RT-qPCR (n=2-4 biological replicates). **C)** *LAMB4* expression during endoderm differentiated from hPSCs. **D)** Schematics of the mesoderm differentiation protocol. **E)** Expression of early mesoderm marker. **F)** Expression of cardiomyocyte-related genes. Previously published RNAseq data was analyzed to assess the expression of laminin chains. **G)** *LAMB4* expression from mesoderm and cardiomyocytes differentiated from hPSCs. Highly expressed laminin genes are shown in green. *LAMB4* shown in red.

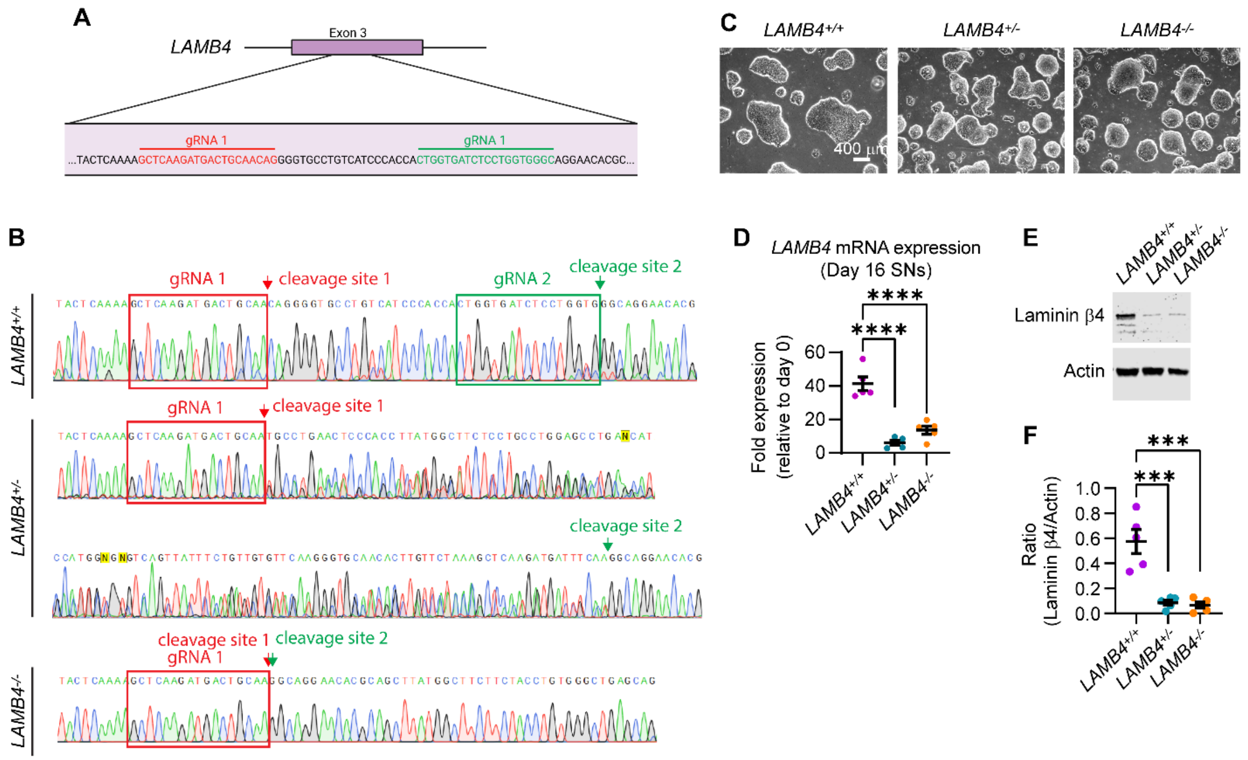


**Figure S3. *LAMB4* editing strategy and characterization. A)** Schematics of the strategy used to edit *LAMB4* by CRISPR/Cas9. **B)** Sequencing results of *LAMB4*$^{+/+}$, *LAMB4*$^{+/-}$, and *LAMB4*$^{-/-}$ hPSCs. **C)** Characterization of *LAMB4*$^{+/+}$, *LAMB4*$^{+/-}$, and *LAMB4*$^{-/-}$ hPSCs by brightfield microscopy. **D)** *LAMB4* expression in SNs differentiated from *LAMB4*$^{+/+}$, *LAMB4*$^{+/-}$, and *LAMB4*$^{-/-}$ hPSCs**.** RNA was isolated on day 16 and *LAMB4* expression was measured by RT-qPCR (n=5 biological replicates). **E)** Laminin β4 levels in *LAMB4*$^{+/+}$, *LAMB4*$^{+/-}$, and *LAMB4*$^{-/-}$ SNs. Total protein of day 20 SNs was immunoblotted for laminin β4 and actin. **F)** Measuring of signal intensity of the immunoblots from **F)** (n=5 biological replicates). For **D)** and **F),** one-way ANOVA followed by Tukey's multiple comparisons test. ***p<0.001, ****p<0.0001. Graphs show mean ± SEM.

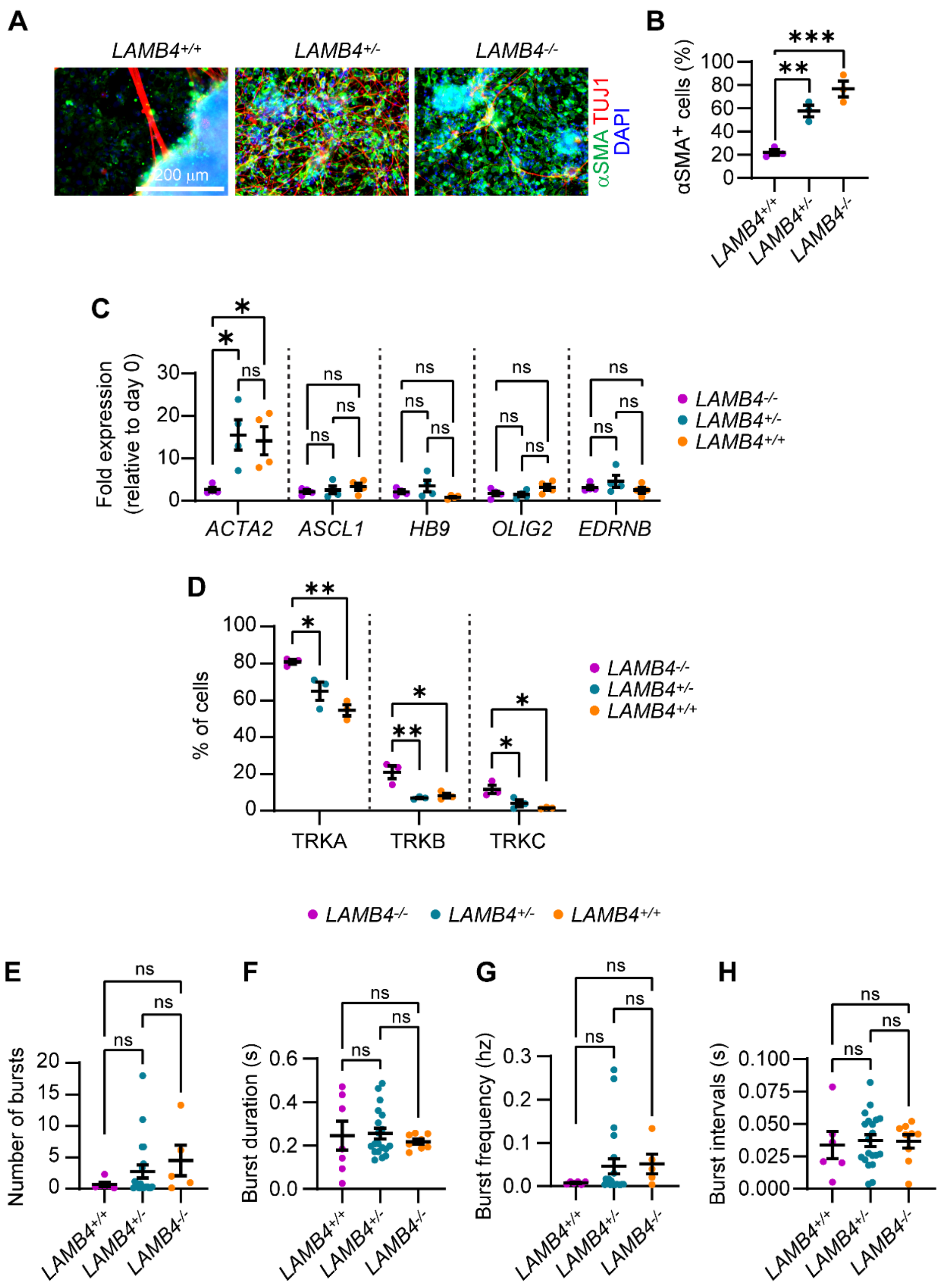


**Figure S4. Characterization of *LAMB4* mutant SNs. A)** Expression of markers upon loss of *LAMB4*. *LAMB4*$^{+/+}$, *LAMB4*$^{+/-}$, and *LAMB4*$^{-/-}$ SNs were fixed on day 20 and stained for the non-neural ectoderm marker α-Smooth Muscle Actin (αSMA). Nuclei were stained with DAPI. **B)**

Percentage of αSMA$^{+}$ cells from **A)** over DAPI was plotted. **C)** Expression of genes not expressed in SNs. RNA of day 20 SNs differentiated from *LAMB4*$^{+/+}$, *LAMB4*$^{+/-}$, and *LAMB4*$^{-/-}$ hPSCs was isolated. Gene expression was measured by RT-qPCR (n=4 biological replicates). **D)** Percentage of cells expressing TRK proteins. Day 20 SNs were stained with TRKA, TRKB, and TRKC fluorescently-tagged antibodies and analyzed by flow cytometry (n=3 biological replicates). **E-H)** Measurement of electrical activity of *LAMB4*$^{+/+}$, *LAMB4*$^{+/-}$, and *LAMB4*$^{-/-}$ SNs. **E)** Number, **F)** duration, **G)** Frequency, and **H)** intervals of bursts were measured. Each dot represents the mean firing rate of 6 wells measured over 40 days (n=4 biological replicates). For **C)**, one-way ANOVA followed by Dunnett's multiple comparisons test. For **B)**, **D)**, **E)**, **F)**, **G),** and **H)** one-way ANOVA followed by Tukey's multiple comparisons test. ns, non-significant, *p<0.05, **p<0.005. Graphs show mean ± SEM.

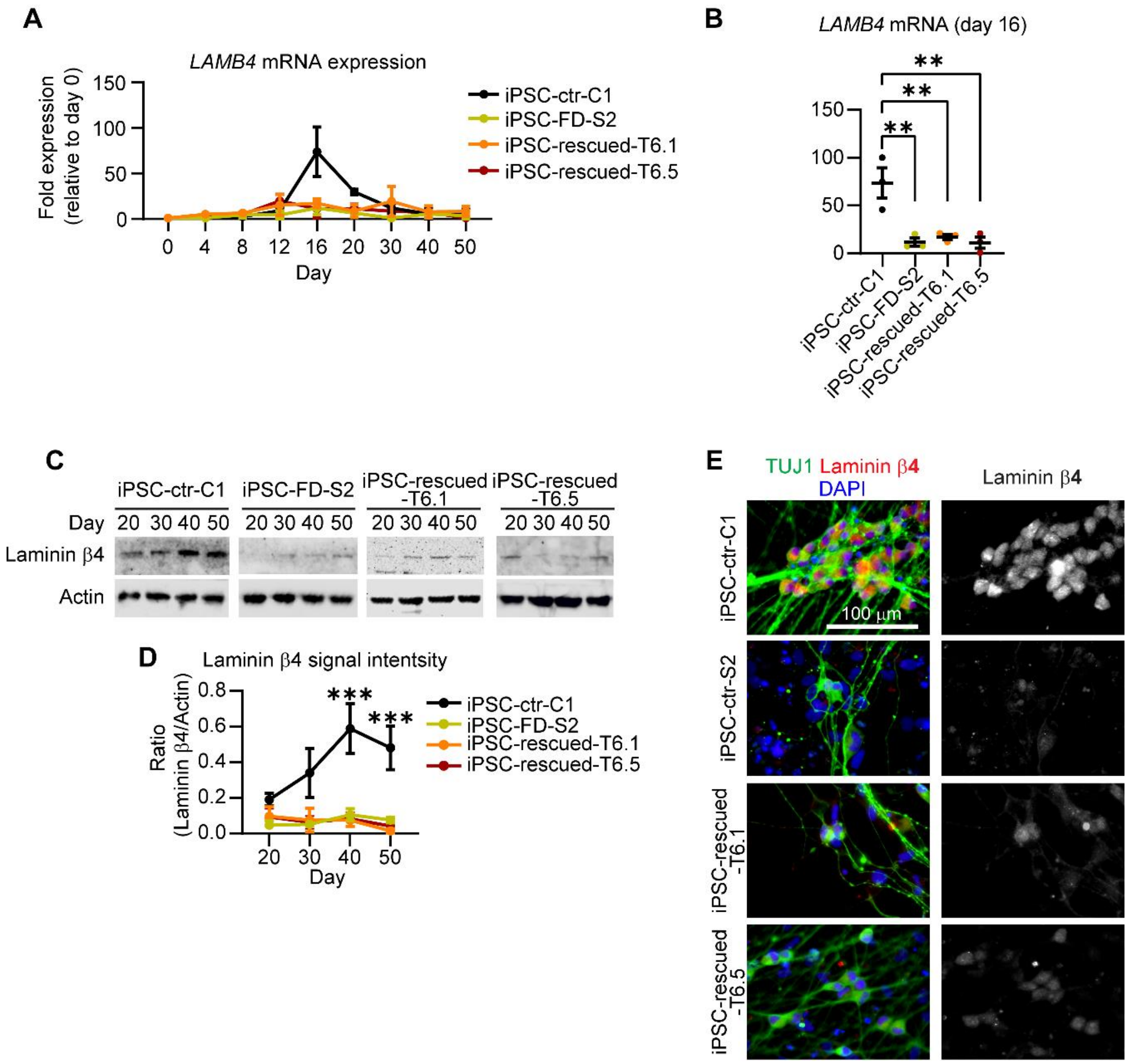


**Figure S5. *LAMB4* expression is not dependent on *ELP1*. A)** *LAMB4* expression in SNs differentiated from *ELP1*-rescued iPSCs of severe FD patients. *ELP1*$^{+/-}$ severe FD iPSCs (iPSC-rescued-T6.1 and iPSC-rescued-T6.5), one severe FD iPSC line (S2), and one healthy iPSC control line (C1) were differentiated into SNs. Total RNA was isolated at the indicated times and gene expression was measured by RT-qPCR (n=3 biological replicates). **B)** *LAMB4* expression by SNs on day 16 shown in **A)** is shown (n=3 biological replicates). **C)** Laminin β4 expression during SN development. iPSC-rescued-T6.1, iPSC-rescued-T6.5, iPSC-FD-S2, and iPSC-ctr-C1 cells were differentiated into SNs. Lysates were collected on the indicated days and were immunoblotted for laminin β4 and actin. **D)** Quantification of signal intensity of immunoblots shown in **C)** (n=3 biological replicates). **E)** Laminin β4 in *ELP1*-rescued severe FD SNs. iPSC-rescued-T6.1, iPSC-rescued-T6.5, iPSC-FD-S2, and iPSC-ctr-C1 hPSCs were differentiated into SNs. Cells were fixed on day 20 and stained for laminin β4, TUJ1, and DAPI. For **B),** one-way ANOVA followed by Tukey's multiple comparisons test. For **D)**, two-way ANOVA followed by Šídák's multiple comparisons test. **p<0.005, ***p<0.001. Graphs show mean ± SEM.

**Supplementary Table 1 -** List of primers used in this study

| Gene | Forward sequence | Reverse sequence |
|---|---|---|
| *SOX10* | CCAGGCCCACTACAAGAGC | CTCTGGCCTGAGGGGTGC |
| *P75NTR* | CCTCATCCCTGTCTATTGCTCC | GTTGGCTCCTTGCTTGTTCTGC |
| *NGN1* | GCCTCCGAAGACTTCACCTACC | GGAAAGTAACAGTGTCTACAAAGG |
| *NGN2* | CAAGCTCACCAAGATCGAGACC | AGCAACACTGCCTCGGAGAAGA |
| *BRN3A* | AGTACCCGTCGCTGCACTCCA | TTGCCCTGGGACACGGCGATG |
| *RUNX1* | CCACCTACCACAGAGCCATCAA | TTCACTGAGCCGCTCGGAAAAG |
| *RUNX3* | GGCAATGACGAGAACTACTCCG | GATGGTCAGGGTGAAACTCTTCC |
| *TRKA* | CACTAACAGCACATCTGGAGACC | ACAGTCAGCTCAAGCCAGACAC |
| *TRKB* | ACAGTCAGCTCAAGCCAGACAC | GTCCTGCTCAGGACAGAGGTTA |
| *TRKC* | CCGACACTGTGGTCATTGGCAT | CAGTTCTCGCTTCAGCACGATG |
| *TBXT* | CCTTCAGCAAAGTCAAGCTCACC | TGAACTGGGTCTCAGGGAAGCA |
| *TBX6* | TCATCTCCGTGACAGCCTACCA | CCGCAGTTTCCTCTTCACACGG |
| *FOXF1* | CAGCCTCACATCACGCAAGG | AGCCGAGCTGCAAGGCATC |
| *TNNT2* | AAGAGGCAGACTGAGCGGGAAA | AGATGCTCTGCCACAGCTCCTT |
| *NKK2-5* | AAGTGTGCGTCTGCCTTTCCCG | TTGTCCGCCTCTGTCTTCTCCA |
| GAPDH | GTCTCCTCTGACTTCAACAGCG | ACCACCCTGTTGCTGTAGCCAA |
| *LAMA1* | GAAGGTGACTGGCTCAGCAAGT | AGGCGTCACAACGGAAATCGTG |
| *LAMA2* | GGCAATCTGAATACACTCGTGAC | TGTGTTGGTCCTCTCAGCATCC |
| *LAMA3* | TAGAGGAAGCCTCTGACACAGG | CCGATAGTATCCAGGGCTACAAC |
| *LAMA4* | GAGATGACTCTCTGCTGGACCT | AGTTCCAGGCAGCCAACAAAGC |
| *LAMA5* | AACCAGATGAGCATCACATTCCTG | ACAGTGTTGCGCGTCTCCGTAT |
| *LAMB1* | GAGGTGTCTCAAGTGCCTGTAC | ACTGGCAGTCAGAGCCGTTACA |
| *LAMB2* | GCGGACTTGTTCTGAGTGCCAA | ACCTGTGAAGCGGTGACACTGA |
| *LAMB3* | GTCACAGAGCAGGAGGTGGCT | GCTTCTGTCAAGACTCTCCAGG |
| *LAMB4* | GTGGAGGCTTTACAACTGGCAG | GGATCATCTGGACACAGGCAAG |
| *LAMC1* | CTGTGAGGTCAACCACTTTGGG | AGCCTTCTCTGCATTCACAGCG |
| *LAMC2* | TACAGAGCTGGAAGGCAGGATG | GTTCTCTTGGCTCCTCACCTTG |
| *LAMC3* | CTGTAACCAGCATGGCACCTGT | ACCTGGCAAACAGCGTTCACAG |
